# Vorinostat Can Reduce an Ongoing Fibrosis and Preserve Key Liver Metabolic Functions by Selectively Altering Mesenchymal Cell Differentiation

**DOI:** 10.64898/2026.11.02.763750

**Authors:** Yuan Guan, Gary Peltz

## Abstract

Liver fibrosis is a major global cause of mortality that is driven by the activation of hepatic stellate cells (HSCs) into extracellular matrix (ECM)-producing myofibroblasts (MyoFs). The lack of predictive human *in vitro* models has severely hampered the discovery of targeted anti-fibrotic therapies. Here, we describe a human iPSC-derived hepatic micro-organoid (microHO) model, which labels collagen producing cells, to study liver fibrosis pathophysiology and screen drug candidates. We discovered that while multiple drugs prevented fibrosis when co-administered prophylactically with TGFβ1, only the FDA-approved pan-HDAC inhibitor Vorinostat (SAHA) successfully reduced an established, ongoing fibrosis and preserved life-sustaining liver metabolic functions in a fibrotic microenvironment, including ammonia detoxification and CYP3A4-mediated drug metabolism. To uncover its mechanism of action, we utilized single-cell RNA sequencing and CRISPR-Cas12a lineage tracing. We found that Vorinostat actively redirects HSC differentiation away from pro-fibrotic MyoFs and toward a distinct, fibrosis-suppressing mesenchymal inhibitory (Mes^Inh^) cell. By reversing TGFβ1-induced epigenetic silencing, Vorinostat opens condensed chromatin to rescue metabolic gene expression and generates anti-fibrotic effectors that neutralize ongoing TGFβ1 signaling. Given its distinct capacity to remodel mesenchymal cell fate and reverse epigenetic repression of metabolic genes at clinically achievable concentrations, Vorinostat represents a highly promising drug for repurposing as a therapeutic for advanced human liver fibrosis.

## Background

Liver fibrosis is a major cause of death (∼1M per year) (*1*) that is often caused by viral infection, metabolism associated steatohepatitis (**MASH**), or chronic alcohol use (*2, 3*). However, extracellular matrix (**ECM**) accumulation after chronic liver injury drives all forms of liver fibrosis (*2, 4*). The key cell is the myofibroblast (**MyoF**), which is generated when fibrogenic stimuli (usually TGFβ1 or PDGF (*5*)) cause hepatic stellate cells (**HSC**) to become the MyoFs that produce excess fibril-forming collagens and ECM proteins (*5–10*). Despite the large patient population, the standard of care for managing liver fibrosis relies upon condition-specific treatments to eliminate the underlying etiology. As examples, eradicating Hepatitis C (*11*) or suppressing Hepatitis B (*12*) with antiviral therapies can reverse an existing liver fibrosis (*13*); and two drugs that both target metabolic drivers - a thyroid hormone receptor-β agonist (resmetirom) (*14–16*) and a GLP-1 receptor antagonist (Wegovy) (*17*) - have received FDA approval for treatment of noncirrhotic MASH with liver fibrosis. There are no universal treatment options for reducing liver fibrosis, and no approved therapy directly targets the HSC activation and MyoF differentiation machinery. The poor quality of available model systems has been a major barrier to identifying new therapies (*18*). Prior *in vitro* models did not have the spectrum of cell types that mediate fibrogenesis and could not reproduce its key feature (i.e., thick collagen filament production) (*19*); and conclusions drawn from rodent models are limited by concerns about their mechanistic fidelity with human liver fibrosis (*20*).

We developed a human-based model system to study the pathophysiology of liver fibrosis, and to identify and characterize the mechanism of action (MOA) of antifibrotic drugs. A human iPSC line with a reporter system that labels collagen producing cells is used to produce a multi-lineage human hepatic organoid grown in microwells (i.e., **microHOs**). MicroHOs have liver lobule architecture with hepatocytes, cholangiocytes around the lumen of functioning bile ducts, endothelial cells, and mesenchymal cells (*18, 21–23*). A fibrotic stimulus causes microHOs to develop a fibrosis with thick collagen fibers throughout the entire organoid, which resembles those in human fibrotic livers (*23*). Transcriptomic comparisons show that cells in microHOs resemble human HSC and MyoF, and MyoF abundance dramatically increases after TGFβ1 exposure (*22, 23*). Five drugs when co-administered with TGFβ1 exhibited anti-fibrotic activity and promoted liver cell regeneration in microHOs (*22, 23*): GSK3β (CHIR-99021, aka GSK3βi), p38 inhibitor (SB202190), Bromodomain 4 inhibitor (ZEN-3694, AKA BRD4i) (*24, 25*), TGFBR1i (A83-01), and a histone deacetylase inhibitor (SAHA, aka HDACi). Vorinostat (SAHA) is an FDA-approved anti-cancer drug that is a pan-histone HDACi (*26, 27*). Here, microHOs are used to further characterize their pharmacodynamic effects. We found that only Vorinostat could both reduce an ongoing fibrosis and preserve essential life sustaining liver metabolic functions; and that its anti-fibrotic MOA was mediated by promoting the generation of mesenchymal cells with potent anti-fibrotic properties.

## Results

### SAHA reduces an ongoing liver fibrosis

To determine if a drug could reduce an ongoing liver fibrosis, microHOs were cultured for 21 days to reach a mature and stable state; TGFβ1 (50 ng/ml) was added on day 21, and test drugs were added one week later (day 28). The extent of fibrosis was serially measured by real time imaging of the Clover signal through day 35 when flow cytometry (FCM) was performed to quantify the shift in cellular populations caused by TGFβ1 ± drug treatments (**Fig. 1A**). As expected, TGFβ1 induced a significant increase in fibrosis from days 25 to 38 (p<0.001), which indicates that there was an ongoing fibrosis when the test drugs were administered on day 28. The fibrosis was markedly reduced (P<0.0001) by the TGFRB1i (A83-01, 10 uM) (p<0.0001), and SAHA (1 uM) also induced significant fibrosis reductions on days 31 (p<0.01) and 38 (p<0.0001) (**Fig.1B-C**). In contrast, the BRD4i (ZEN-3694 2 uM), which reduced fibrosis when co-administered with TGFβ1 (*22*), did not significantly reduce the ongoing fibrosis, but fibrosis was reduced by the p38i (SB202190, 10 uM) (P<0.001) and GSK3βi (CHIR-99021, 3 uM) (p<0.001) (Fig. 1B). Analysis of microHO size indicated that drug toxicity was not responsible for the antifibrotic effects (**Fig. S1**). Moreover, Trichrome staining confirmed that SAHA decreased the amount of ECM in TGF*β*1-treated microHOs (**Fig. 1D**). Bile acid production was also inhibited in TGFβ1-treated microHOs, but bile production was increased after SAHA treatment (**Fig. 1E**). FCM scatter (**Fig. 1F**) and boxplots (**Fig. 1G**) quantitatively assessed the effects of TGFBR1i or SAHA on fibrosis and liver cell regeneration on day 38. Since Epithelial Cell Adhesion Molecule (EPCAM) is a hepatocyte and cholangiocyte marker, an increase in EPCAM^+^ cells indicates that liver tissue is regenerating. TGFβ1 increased the number of Clover^+^ cells and decreased EPCAM^+^ cells in microHOs. TGFRB1i or SAHA treatment markedly decreased the Clover^+^ population, which confirms their anti-fibrotic effect, but also increased the EPCAM^+^ population (Fig.1F-G). These results indicate that the liver cells in microHOs can regenerate after TGFRB1i or SAHA treatment in the presence of an ongoing fibrosis.

**Figure 1.**
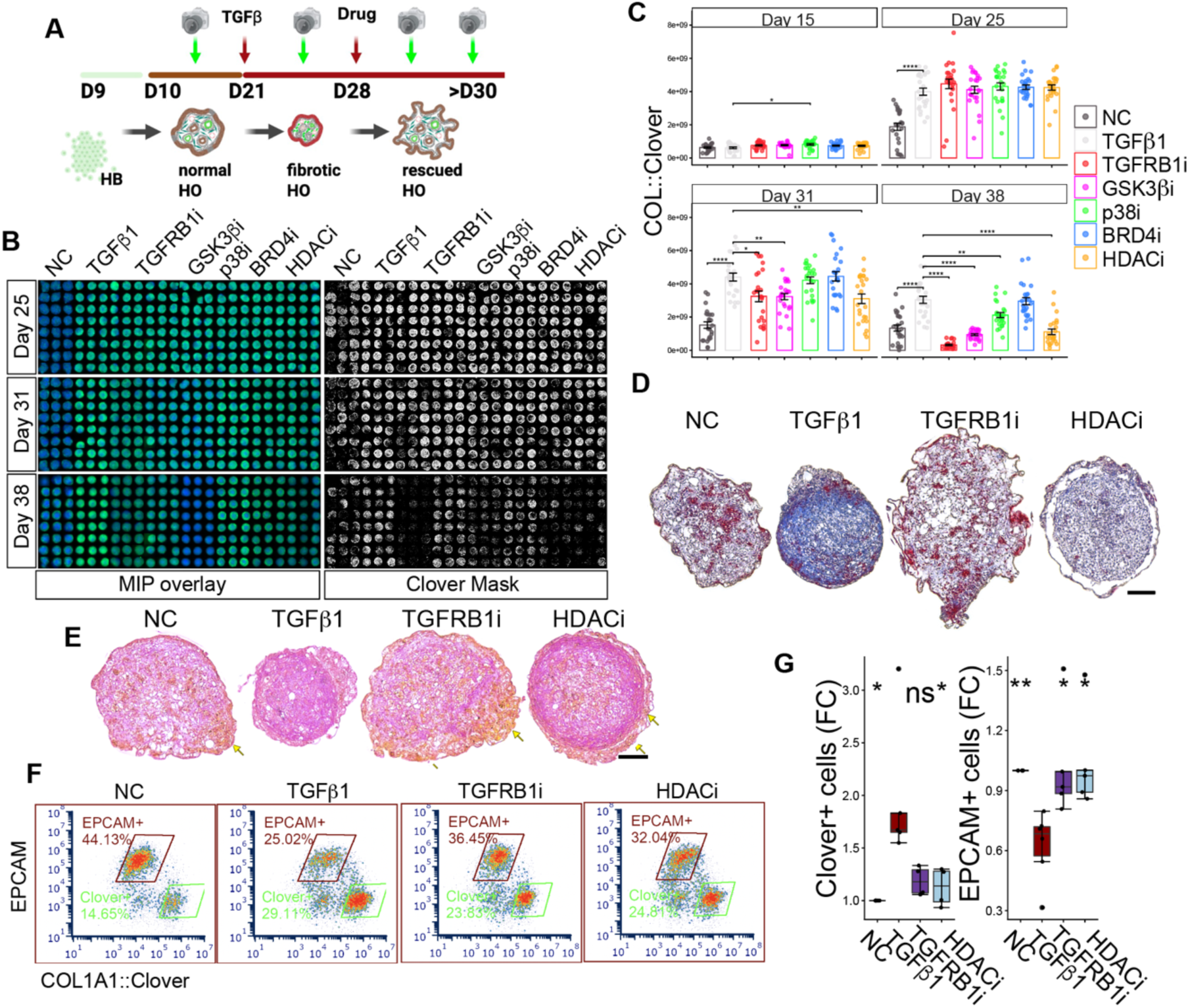
An HDAC inhibitor SAHA (Vorinostat) reduces an ongoing fibrosis in microHOs and promotes epithelial cell regeneration. (**A**) A diagram showing how a fibrosis in day 21 mature microHOs is induced by TGFβ1 (50 ng/ml) addition. These drugs were administered one week later (day 28) while the fibrosis was ongoing: SAHA (1 uM); BRD4i (ZEN-3694 2 uM); TGFBR1 inhibitor (A83-01, 10 uM); GSK3βi (CHIR-99021, 3 uM); and p38i (SB202190, 10 uM). The extent of fibrosis is serially assessed through day 38 by measuring the COL1A1::Clover signal at the indicated timepoints, and flow cytometry (FCM) analysis is performed on day 38 to assess the effect of the drugs and TGFβ1 on the cellular composition of the microHOs. (**B**) Representative images of day 38 microHOs in microwells that were treated as described in (A) with an overlay for detecting nuclear Hoechst 33343 staining (MIP) or the COL1A1::Clover signal (Clover Mask). (**C**) The COL1A1::Clover green fluorescence signal is directly linked to the production of Type I Collagen, which generates an intracellular fluorescent signal after cleavage of the COL1A1: Clover peptide. Each treatment and time point are the average of measurements made on >30 microHOs. For each drug + TGFβ1 treatment: *, p-value<0.05; **, p-value<0.01; ***, p-value<0.001; and ****p-value<0.0001 vs TGFβ1 alone. TGFβ 1induces a significant increase in fibrosis that is significantly reduced by treatment with SAHA on day 28. The BRD4i had a small effect on fibrosis, and SAHA had a greater effect on the ongoing fibrosis than the GSK3βi and p38i. The TGFBR1i served as a positive control that substantially reduced the ongoing fibrosis. (**D**) Images of Trichrome-stained day 38 microHOs treated as described in (A). The TGF*β*1-treated microHOs show a marked increase in collagen-rich connective tissue (blue-stained regions) relative to NC microHOs, which only have a thin layer of connective tissue. The TGF*β*1-induced increased in collagen was markedly reversed by addition of TGFBR1i or HDACi. (**E**) Representative images of day 38 microHOs treated as described in (A) that were stained to detect bile acids (Hall’s method). TGFβ1 markedly decreases bile acid production in microHOs, while bile production (yellow arrows) is recovered after treatment with TGFBR1i or HDACi. All scale bars are 100 μm. (**F, G**) Representative FCM scatter and box plots show the increase in Clover^+^ cells (fibrosis) and the decrease the EPCAM^+^ population in TGFβ-treated microHOs. EPCAM is a marker for liver parenchymal cells (hepatocytes and cholangiocytes), which provides an indication of whether parenchymal cells are regenerating. SAHA administration decreased Clover^+^ cells and increased EPCAM^+^ cells in the TGFβ-treated microHOs. The boxplots show the fold-change (FC) in Clover^+^ and EPCAM^+^ populations as measured by FCM in normal control (NC), TGFβ (T), TGFβ+drug (either TGFBR1i (Ti) or SAHA)-treated microHOs. Each group has 4 experimental replicates. The p-values for the TGFβ-treated microHOs were calculated (ns, not significant; *, p-value<0.05; *, p-value<0.001) ** relative to NC; and p-values for the TGFβ + drug treated microHOs were calculated relative to TGFβ-treated microHOs. TGFβ1 caused a significant increase in Clover^+^ cells and a significant decrease in EPCAM^+^ cells. However, TGRBR1i and SAHA caused a marked decrease in Clover^+^ cells and an increase in EPCAM^+^ cells, which were not changed relative to NC microHOs.

### SAHA maintains essential liver metabolic functions

We investigated whether these antifibrotic drugs could maintain liver metabolic functions that are essential for human health. For example, hepatic encephalopathy (cognitive impairment, lethargy, and confusion that can progress to comma) is a major liver failure complication. It develops when liver urea cycle enzymes are unable to metabolize the ammonia (*28–30*), which is continuously generated in the gastrointestinal tract and transported via blood to liver, to a water-soluble compound (urea) that is excreted by the kidney. An ammonia buildup is harmful because the brain converts ammonia to glutamine, which causes osmotic swelling, mitochondrial toxicity, and neurotransmitter imbalance (*30–32*). Therefore, the effect of TGFβ1, and of four anti-fibrotic drugs when co-administered with TGFβ1 on day 13, on the level of expression of urea cycle enzyme mRNAs (*CPS1, OTC, ASS1, ASL, ARG1* and *NAGS*) in day 21 microHOs was examined (**Fig. 2A**). TGFβ1 markedly decreased urea cycle mRNA expression (vs control), while SAHA (1 uM) or TGFBR1i (10 uM) co-administration rescued urea cycle enzyme mRNA expression. Despite their anti-fibrotic effect, GSK3βi (3 uM) and p38i (10 uM) could not reverse this TGFβ1 effect (**Fig. 2B**). Urea production by microHOs was measured to further evaluate SAHA’s effect. As anticipated, TGFβ1-induced fibrosis significantly reduced urea secretion (p < 0.0001 vs NC); while TGFBR1i (10 uM) (p<0.0001) or SAHA (1 uM) (p<0.001) co-administration protected against the TGFβ1-induced decline in urea generation (**Fig. 2C**). TGFBR1i (p = 3.1×10^-13^, vs TGFβ) or SAHA ( p=0.00081) maintained urea secretion at control microHO levels. Thus, SAHA co-administration can preserve the liver’s ability to detoxify ammonia in the presence of a fibrotic stress.

**Figure 2.**
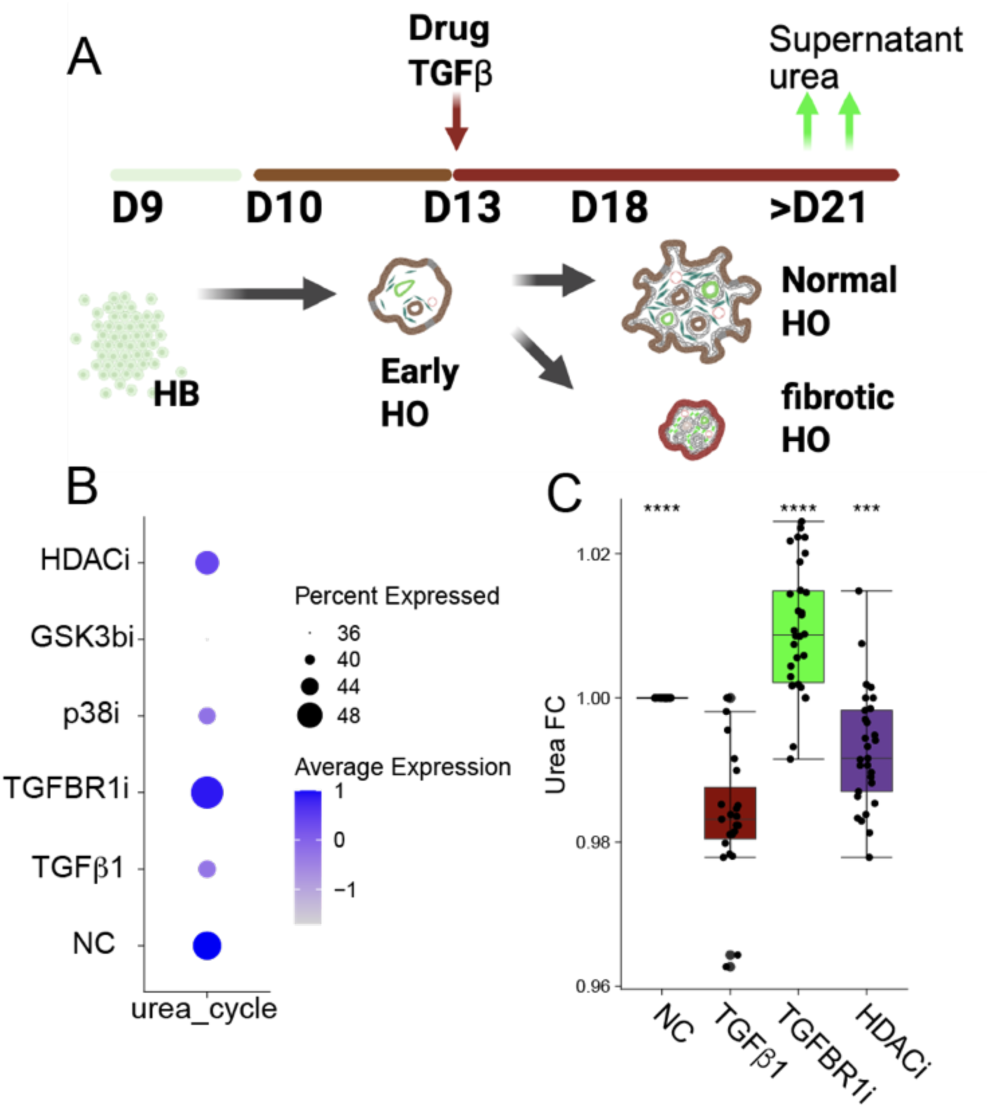
SAHA preserves the ability of microHOs to metabolize ammonia to urea. (**A**) The microHO platform used to assess TGFβ1 and drug effect on urea production. COL1A1::Clover iPSCs are differentiated into hepatoblasts (HB) on day 9, and 10,000 HBs are placed in each microwell on day 10. TGFβ1 (50 ng/ml) ± drug are added to each well on day 13, and the cultures are further differentiated into hepatic organoids. Samples for scRNA-Seq and urea measurements are collected on day 21. (**B**) SAHA treatment increases urea cycle mRNAs in fibrotic microHOs. microHOs were treated with TGFβ (50 ng/ml) and the following drugs on day 13 (as in (A)): HDACi (SAHA 1 uM); TGFBR1 inhibitor (A83-01, 10 uM); GSK3βi (CHIR-99021, 3 uM); and p38i (SB202190, 10 uM). Module scores were calculated from the scRNA-Seq data to assess the relationship between the transcriptomes of normal control (NC), TGFβ1 and TGFβ+drug-treated microHOs for the urea cycle core gene set (*CPS1, OTC, ASS1, ASL, ARG1* and *NAGS*). The size and color of each dot represents the percentage of cells expressing urea cycle enzymes and their average expression level. TGFβ1 markedly decreases urea cycle mRNA expression (vs NC), while SAHA or TGFBR1i co-administration restores urea cycle enzyme expression in fibrotic microHOs. In contrast, the GSK3βI or p38i could not reverse the TGFβ1 effect on this critical liver function. (**C**) This boxplot shows the fold change (FC) in the level of urea secretion by microHOs after four different experimental treatments: NC (baseline set to FC = 1.00), fibrotic microHOs treated with TGFβ1 on day 13 (TGFβ), TGFβ1 and TGFBR1i (10 uM) treated microHOs, or TGFβ1 + SAHA (1 uM)-treated microHOs. To evaluate ammonia conversion capacity, microHO culture supernatants (50 ul per organoid, >16 per group) were harvested on day 21, and urea production was measured. The central horizontal line represents the median, the boxes represent the interquartile range (IQR), and whiskers extend to the minimum and maximum values. The individual black dots represent independent replicates. Statistical significance was assessed using a student’s t-test *** p < 0.001, **** p < 0.0001. The TGFβ1-treated microHOs were compared with NC, and all TGFβ1 + drug groups were compared to TGFβ1-treated microHOs.

Cytochrome P450 3A4 (CYP3A4) is the most abundant drug-metabolizing enzyme in human liver; it is responsible for the metabolic clearance of ∼50% of all prescription drugs (*33, 34*). TGFβ rapidly downregulates CYP3A4 activity by decreasing the expression of the primary transcriptional regulator (Pregnane X Receptor, PXR) of *CYP3A4* in hepatocytes (*35*), which is further decreased by the liver fibrosis microenvironment (*36, 37*). Ketoconazole blocks basal and inducible *CYP3A4* expression by acting as a PXR antagonist (*38*) and directly inhibits CYP3A4 enzyme activity by binding to its active site (*39, 40*). Therefore, CYP3A4 activity was measured in control and TGFβ1-treated microHOs in the presence or absence of ketoconazole or rifampin. After 6 hours, CYP3A4 enzyme activity was not statistically different between control, TGFβ1-, or TGFβ1+drug-treated microHOs, irrespective of whether a CYP3A4 inducer or inhibitor was present. There was a minimal level of CYP3A4 activity in the presence of the CYP3A4 inhibitor (Ketoconazole) in all groups at the 6 and 24-hour time points, which confirms assay specificity for CY3PA4 activity (**Fig. 3**). However, notable differences appeared after 24 hours of exposure to the CYP3A4 inducer (Rifampicin). TGFβ1 induced a significant decrease in rifampin-induced CYP3A4 activity within 24 hours in microHOs (p<0.05 vs controls) (Fig. 3), which reflects changes noted in fibrotic human liver tissue. While the TGFB1i did not block the TGFβ1-induced decrease in CYP3A4 activity, SAHA co-administration significantly increased CYP3A4 activity in TGFβ1-treated microHOs to above the control level (p<0.05) (Fig. 3). Hence, besides protecting against TGFβ1-induced fibrosis, SAHA preserves critical hepatocyte metabolic functions, which include ammonia and drug-metabolism.

**Figure 3.**
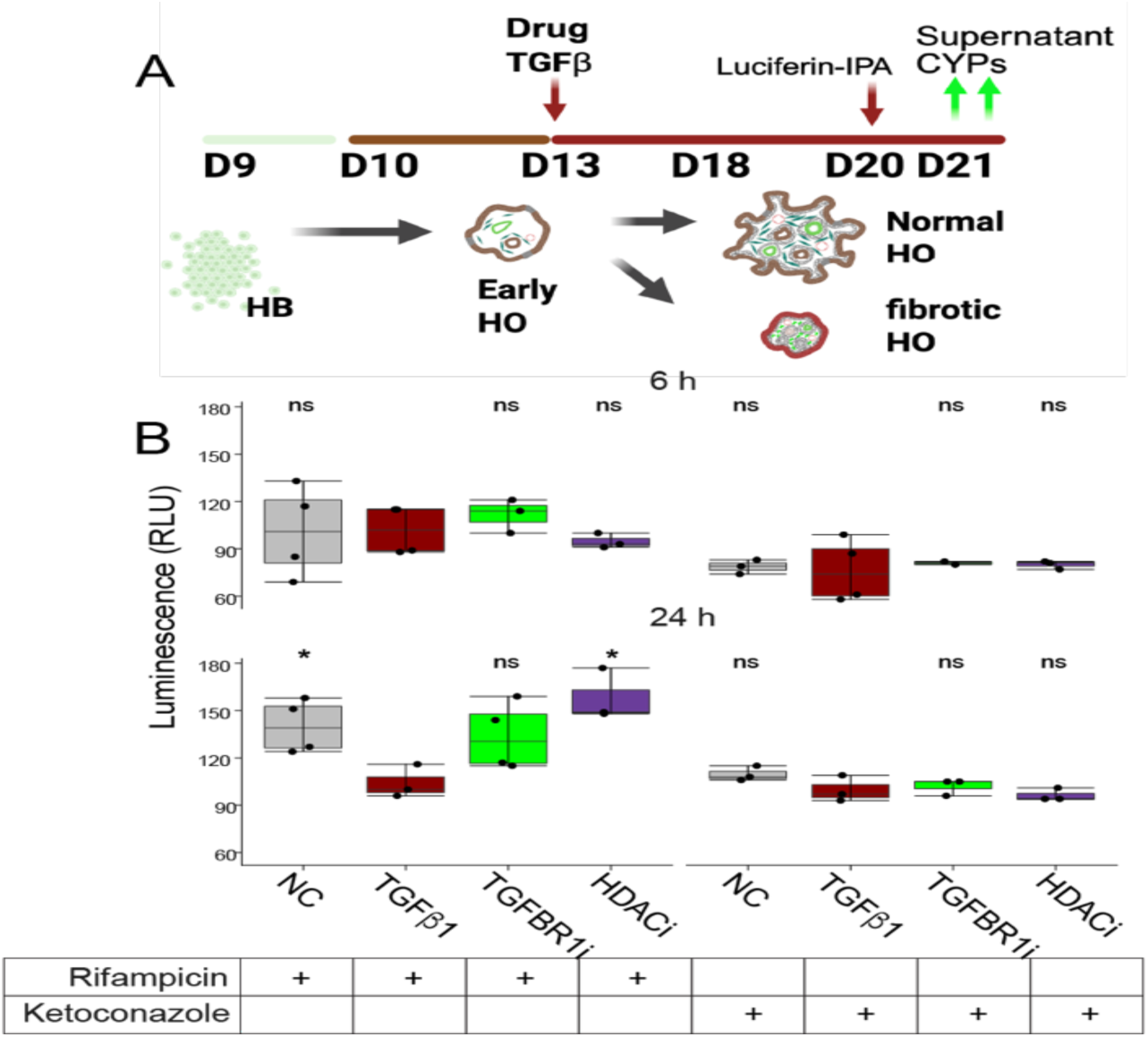
SAHA restores CYP3A4 activity in fibrotic microHOs. microHOs were incubated with normal culture media (NC) or TGFβ1 with or without drugs on day 13. Then, a CYP3A4 inducer (Rifampicin, 25 μM) or inhibitor (Ketoconazole, 1μM) was added on day 21. The following drugs were added to the TGFβ1-treated microHOs: TGFBR1i (A83-01, 10 uM) or HDACi (SAHA, 1 uM). CYP34A activity was measured by quantifying luminescence (Relative Light Units, RLU) after a 6- or 24-hour incubation with a derivative that is converted by CYP3A4 to a luciferase substrate. While the TGFBR1i could not restore rifampicin-mediated CYP3A activity induction in fibrotic microHOs, SAHA restored rifampicin’s ability to induce CYP3A4 activity. Each datapoint is the average and SE of measurements made on 4 different batches (8 per batch) of microHOs. The statistical significance of the 6 or 24-hour datapoints are calculated by comparing TGFβ1-treated fibrotic microHOs vs NC (* p < 0.05; ns = not significant) or by comparing drug + TGFβ1-treated microHOs vs TGFβ1-treated fibrotic microHOs.

### The MOA of SAHA’s anti-fibrotic effect

We coupled scRNA-seq with a CRISPR cell barcode (iCas12-DAISY) lineage tracing system (*22*) to characterize SAHA’s effect on fibrotic microHOs (**Fig. 4A-B**). Single-cell transcriptomes from 350,000 cells in mature vehicle (NC), TGFβ1 (T), TGFβ1 + Ti, and TGFβ1 + SAHAx-treated microHOs. TGFβ1 and drug were co-administered on day 13. UMAPs show that microHO cells in the four treatment arms could be partitioned into 16 distinct clusters, which were differentially affected by the treatments (**Fig. 4C**). TGFβ1 drives cells into regions that are largely absent in vehicle controls; whereas Ti co-treatment largely causes the cells to reoccupy the vehicle territory. SAHA co-administration caused substantial portions of the TGFβ1 cell occupied areas to be vacated, and the vehicle occupied regions were partially re-occupied by cells, which indicates that the non-fibrotic cell distribution is partially recovered. The UMAPs indicate that SAHA has a selective cluster-specific effect rather than a complete phenocopy of TGFβ type 1 receptor blockade. The key Ti or SAHA-induced changes were identified by implementing a two-contrast rule, which compared the changes in the percentage of the 16 annotated cell types in (i) TGFβ1 vs NC microHOs and then in (ii) TGβ1 vs TGFβ1+drug-treated microHOs. This identifies the cell populations whose percentage is altered by TGFβ1, which are then altered by SAHA or Ti co-treatment (**Fig. 4C**). Ti serves as a benchmark indicator for a maximal anti-fibrotic drug effect. Ti robustly reverted all eight of the cell types whose percentages were altered in this manner (Tier A) across all experimental batches (i.e., the percent change in the cell type caused by TGFβ1 at FDR ≤ 0.05 (q₁) was also changed in the TGFβ1+Ti arm at FDR ≤ 0.05 (gated q₂). For example, Ti co-administration virtually eliminated MyoFs (34.57% to 0.69%, rev = 1.20, q2 = 7.2 x 10^-5^), fully restored HSCs to their basal level (3.68% to 22.06%, rev = 0.82), and expanded hepatocytes to above their basal level (1.11% to 19.47%, rev = 2.05) (**Fig. 4D, right; Fig. S2**). TGFβ1 and SAHA jointly altered four cell populations, but the strongest effect of SAHA co-administration was an increase in Mes^Inh^ cells from 8.55% in NC to 20.75% (TGFβ1+SAHA) that was observed in all seven experimental batches (q2 = 0.0011, rev = −2.04). SAHA partially reduced MyoFs from 32.1% to 22.7%, (rev = 0.36). Robustness tests confirmed that SAHA co-administration induced significant changes across all 7 experimental batches only for the Mes^Inh^ and MyoF cells (**Fig. 4D-E, left; Fig. S3**). To further characterize SAHA’s effect on the three key mesenchymal populations (HSC, MyoF, and Mes^Inh^), barcode labelled cells (n=81,863) were mapped onto the 350,000 microHO cell reference atlas, which included the 16 previously annotated cell types (*41*) (**Fig. S4**). This dataset reproduced the previously observed effects of Ti and SAHA co-administration on MyoFs (complete or partial reversion, respectively) (**Fig. S5**). The cross-dataset consensus is that Ti reduces Mes^Inh^ cells to below baseline levels (from 3.5% to 0.8%), while SAHA increases the Mes^Inh^ population to 13.0%. Thus, SAHA’s antifibrotic effect is distinct from that of the Ti and is mediated by its selective effect on mesenchymal cell differentiation, which generates Mes^Inh^ cells.

**Figure 4.**
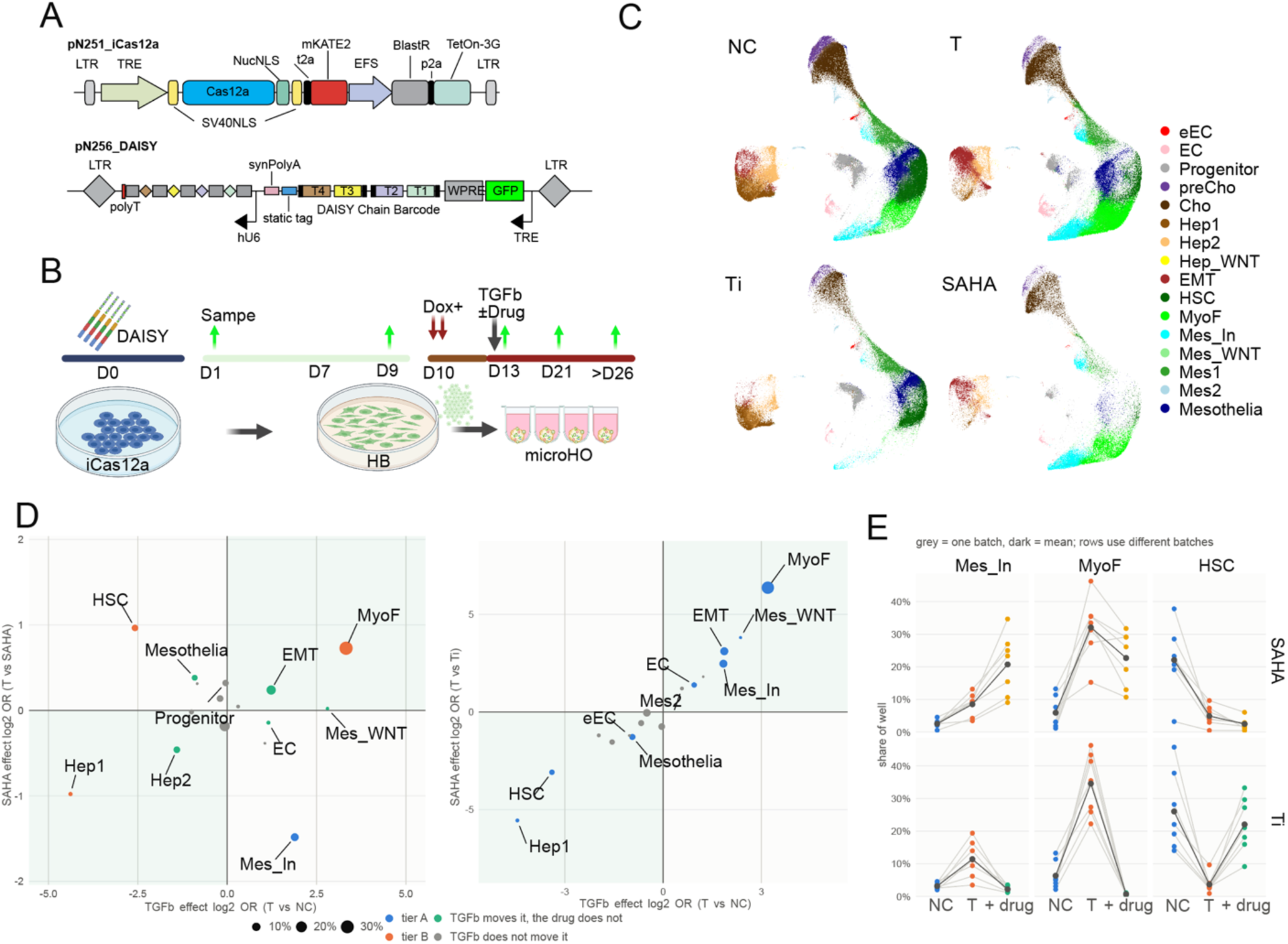
(**A**). The iCas12a and DAISY barcode system. Inducible Cas12a expression is controlled by doxycycline exposure. DAISY is an integrating cassette with a 10 bp integration barcode and four target sites (T1-T4) for CRISPR editing (at cassette positions 45, 61, 105 and 121). (**B**) Cellular barcode and transcriptome analysis for microHO experiments. DAISY barcodes are introduced by lentivirus (MOI of 1∼2) to iPSCs before Day 0. The treatment arms for the microHOs are NC, vehicle; T, TGFβ1 (50 ng/mL); Ti, (TGFBR1i, 10 µM); and SAHA (1 µM). Doxycycline (Dox 500 ng/ml) is added on day 10. TGFβ1 (±drug) is added on Day 13, immediately after the early HO sample is taken; and then sampling occurs on days 20, 21, 26, and 30. Because the recorder cassette is delivered once before Day 0, every mark written prior to Day 13 is shared by all treatment arms, and only marks written after Day 13 are treatment arm-specific. (**C**) A UMAP showing microHO cells in the 4 treatment arms superimposed on the microHO atlas reference. The MyoFs (bright green) increase from 5.8% in NC to 29.9% in TGFβ-treated microHOs, but this increase is blocked by Ti co-administration (0.6 %). In contrast, Mes^Inh^ (cyan) cells increase from 3.5% to 10% in TGFβ-microHOs; Ti co-administration reduces Mes^Inh^ cells to 1.9%; but SAHA co-administration increases Mes^Inh^ cells to 22.1%, which is higher than in TGFβ microHOs. HSC (dark green) decreases from 19.7% in NC to 4.6% in TGFβ-microHOs, but recover (17.7 %) when Ti is co-administered. **(D)** Two-contrast selection of drug-responsive cell types for SAHA (left) and Ti (right). The points represent each of the 16 annotated cell clusters plotted by the TGFβ1-induced change (x-axis: log₂ OR, T vs. NC) against the drug-induced change (y-axis: log₂ OR, T vs. T+drug; n = 7 batches). Point size denotes the cluster share in the T arm; the colors indicate the significance tiers based upon a dual-contrast threshold (FDR ≤ 0.05 for both TGFβ and drug effects). Diagonal quadrants mark phenotypic reversion toward baseline, while off-diagonal quadrants indicate amplification. Mes^Inh^ cells are the only Tier A population for SAHA co-administration, exhibiting significant amplification (q₂ = 0.0011 across all 7 batches), whereas MyoF and HSC qualify as Tier B. Conversely, Ti robustly reverts eight Tier A populations, highlighting SAHA’s cluster-selective efficacy. (**E**) The effect of Ti (Top) or SAHA (Bottom) co-administration on three mesenchymal cell populations in microHOs. The columns indicate the Mes^Inh^, MyoF and HSC cell types; the change in percent of each cell type in NC → T → SAHA microHOs (Top) or between NC → T → Ti microHOs (bottom) are shown. There is one grey line per experimental batch, the dark line is the mean; the third position in each row (+ drug) is the drug indicated at the left of the graph. All six panels share a y-axis to enable cell type effects to be compared.

It was important to characterize the differentiation pathway(s) that connects the three key mesenchymal populations (HSC, MyoF, and Mes^Inh^) in microHOs and to understand SAHA’s effect on them. HSC could differentiate into MyoF that subsequently differentiate into Mes^Inh^ cells, and SAHA’s MOA was to promote the differentiation of MyoF into Mes^Inh^ cells. Alternatively, HSC could differentiate into either MyoF or Mes^Inh^ cells, and SAHA could promote HSC differentiation into Mes^Inh^ cells. To distinguish between these possibilities, we simultaneously analyzed the transcriptome and CRISPR edited barcodes in cells in control, TGFβ1-, TGFβ1 and drug-treated microHOs. For this analysis, the drugs (either Ti or SAHA) were co-administered with TGFβ1 on day 13 (**Fig. 5A**). Claude code (Default model selection with high effort setting) was used to enhance our analysis of the scRNA-seq and barcode data. As described in Supplemental Note 1, we established the lineage recorder fidelity of the DAISY barcode; we demonstrated that Cas12a expression was doxycycline (Dox)-dependent, had negligible background leakage, and identified an optimal Dox induction schedule; and there was a high level of editing diversity, which enabled cell lineage reconstruction (**Fig. S6).** Barcode diversity was preserved throughout long-term microHO culture and progressed throughout microHO differentiation without any technical bottlenecks; which ensured that there was a robust, depth-controlled barcode recording substrate for downstream phylogenetic reconstruction; (**Figs. S6-13**).

**Figure 5.**
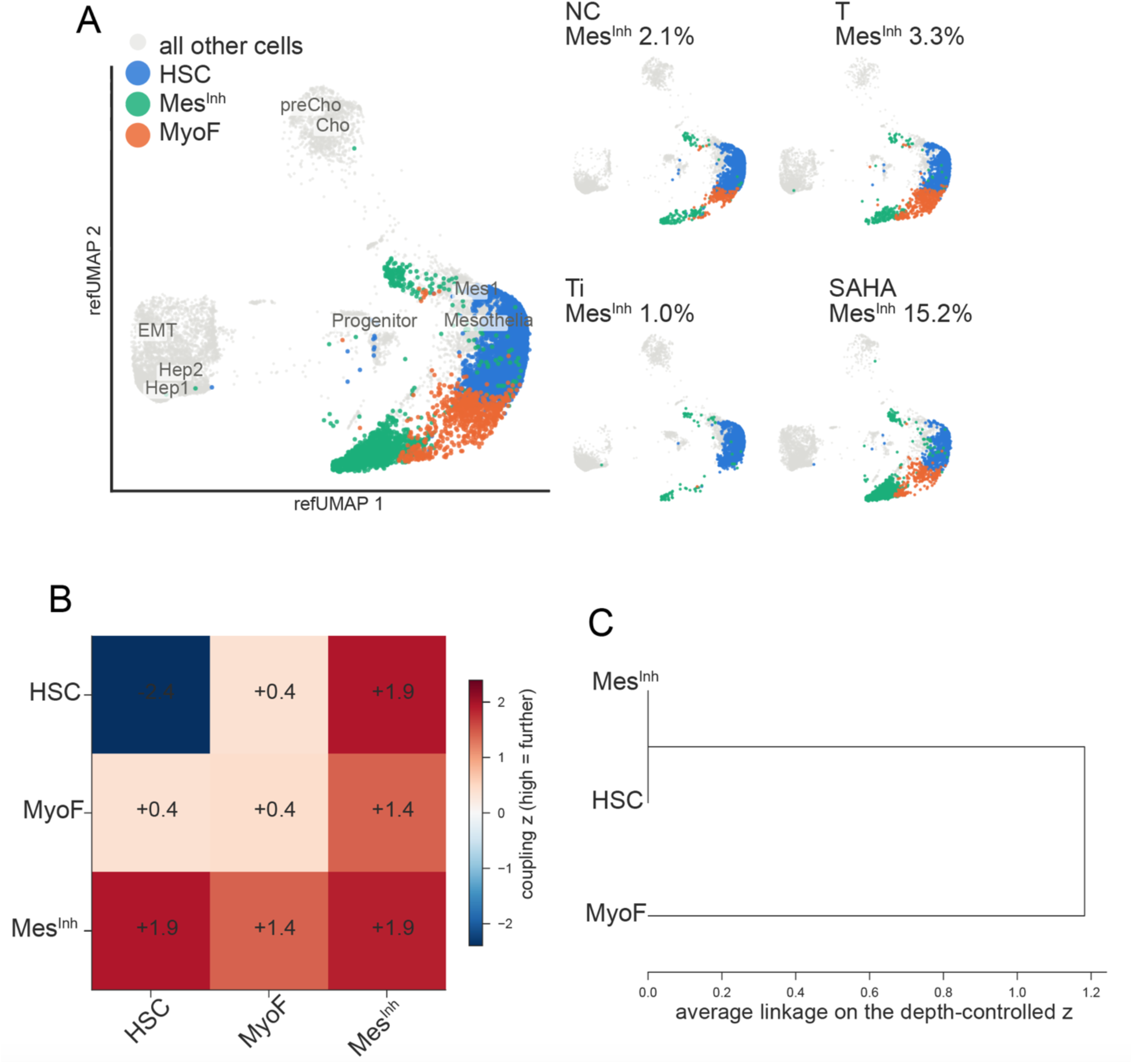
Lineage relationships among HSC, MyoF and Mes^Inh^ cells. (**A**) *Left:* A UMAP shows the three mesenchymal types in day 21 microHOs in color (Mes^Inh^ 1455 cells; MyoF 684 cells; and HSC, 9084 cells) relative to the other cell types (grey background, 16315 cells). *Right:* UMAP showing the Mes^Inh^, MyoF and HSC cells in the 4 treatment arms in color relative to the other cell types (grey background). The marked increase in Mes^Inh^ in the SAHA treated microHOs is readily apparent. **(B)** Pairwise lineage distance matrix among mesenchymal subpopulations. Heatmap depicting evolutionary coupling *z*-scores among HSC, MyoF, and Mes^Inh^ cells derived from clonal barcode analysis. The coupling distances were quantified via cassiopeia.tl.compute_evolutionary_coupling across 200 random permutations/shuffles. The values represent depth- and treatment-arm-controlled normalized coupling scores, where a lower *z*-scores denote significant lineage proximity (closer barcode sharing than by chance) and higher *z*-scores represent greater lineage separation. This matrix is the basis for hierarchical average-linkage tree shown in (C). (**C**) Analysis of cell barcode data obtained from microHOs on days 21 and 26 indicates that HSC can differentiate into Mes^Inh^ or MyoF. Mes^Inh^ and HSC were consistently the closest pair, which was determined by a depth- and arm-controlled calculation of lineage distance on both days and under clone resampling (support 0.78 and 0.70 against 0.33 expected by chance).

Restricting the phylogenetic analysis to three mesenchymal cell types - which included 1,027 cells across 72 clones (43.4% of the tree) at Day 21, and 772 cells across 48 clones (38.2%of the tree) at Day 26 reduced the comparison space and substantially increased the number of per-cell comparisons. This enabled ‘cassiopeia.tl.compute_evolutionary_coupling’, which assesses lineage distances by determining whether the barcodes in two cell groups are closer than by chance (*42–44*), to characterize their phylogeny. Analysis of the (depth- and arm- controlled) distance matrix identified HSC and Mes^Inh^ cells as the most closely related cell pair at both analyzed time points (**Fig. 5B**). Hierarchical average-linkage clustering confirmed that HSC and Mes^Inh^ cells were grouped before either cell type merged with MyoF on both days (bootstrap support of 0.78 and 0.70 on days 21 and 26 vs 0.33 expected by chance) (**Fig. 5C**). Consistent with the barcode results, RNA velocity analysis, which assesses cellular differentiation by analyzing mRNA splicing kinetics (*45*), mapped a developmental path originating from HSC that branches into distinct MyoF and Mes^Inh^ cells (**Fig. 6A**). In control microHOs, HSCs can differentiate into other cell types, but TGFβ1 forces HSCs to predominantly become MyoFs (Fig. 6B-C). Ti co-administration essentially eliminates the MyoF population and completely reverses the TGFβ1 effect. In contrast, SAHA co-administration redirects HSC differentiation to become Mes^Inh^ cells, which were identified by three different cell trajectory analysis methods Velocity Pseudotime (*45*), CytoTRACE2 (*46*), and Monocle 2 Pseudotime (*47*) as the most terminally differentiated of the 3 cell types (Fig. 6E-M and **S14**). Analysis of the differentially expressed genes (DEGs) in Mes^Inh^ cells indicate that that they express multiple proteins that suppress fibrosis in liver and other organs (discussed below). Hence, SAHA induces HSC to become a terminally differentiated mesenchymal cell type (Mes^Inh^) that can suppress fibrosis.

**Figure 6.**
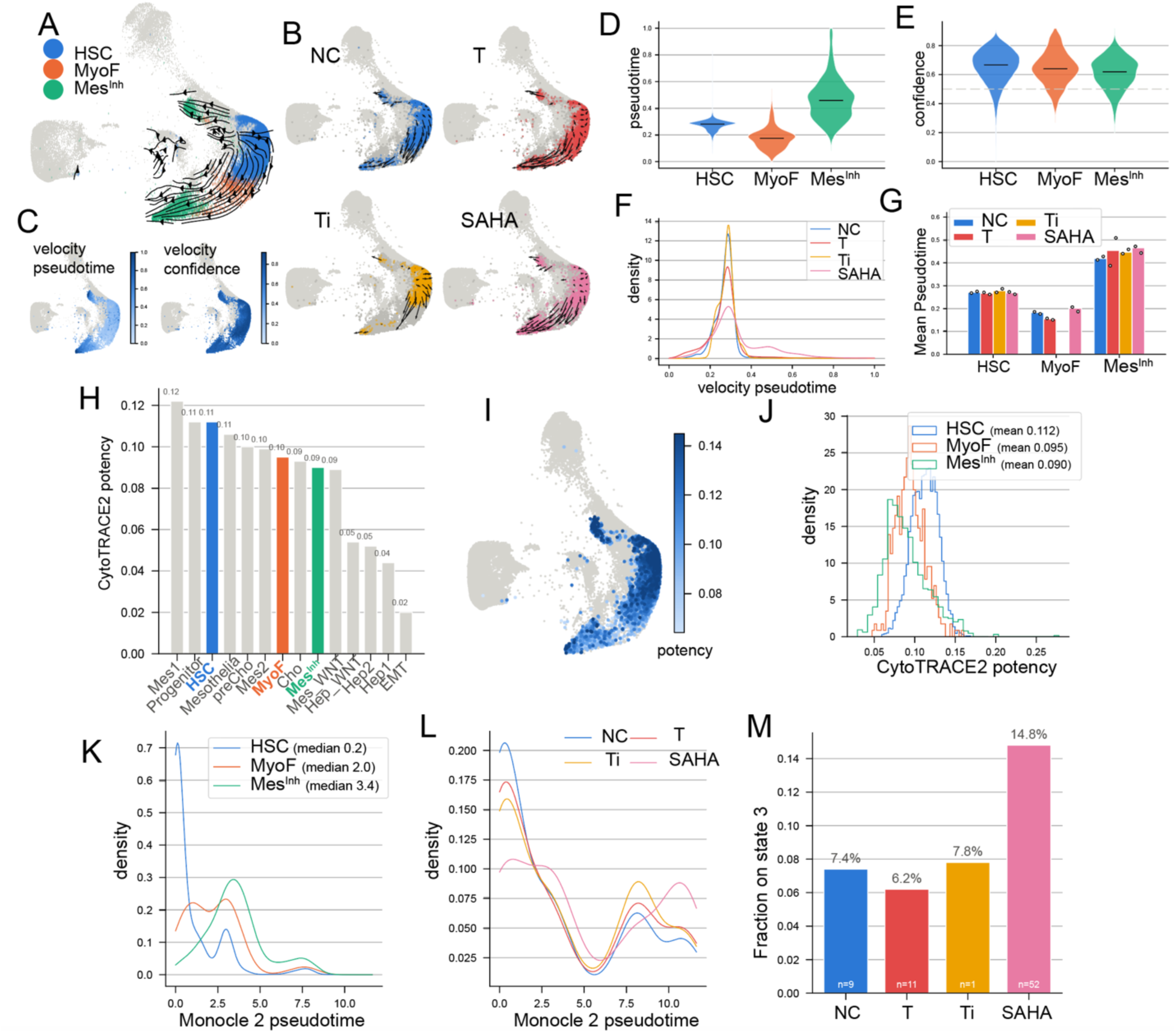
scRNA-seq analyses examine the differentiation state and relationships of HSC, Mes^Inh^ and MyoF cells; and the effect of TGFβ1 and SAHA coadministration on their differentiation. (**A-G**) RNA velocity analyses indicate that Mes^Inh^ are the most differentiated of the 3 cell types; and identifies a developmental path originating from HSC that branches into MyoF and Mes^Inh^ cells. (**A**) The RNA velocity stream is projected onto a reference UMAP, which has 26,063 cells of the three cell types of interest shown in color (HSC blue, MyoF orange, Mes^Inh^, green) against a grey backdrop of 60,000 other cell types. The arrow length is the mean projected velocity of the cells in each grid square. (**B**) The same velocity field is divided into the four treatment arms (NC, 6,408 cells; TGFβ1 (T) 5,195 cells; Ti coadministration 5,467 cells; SAHA coadministration 3,440 cells) with grid-averaged velocity arrows. (**C**). The velocity pseudotime (left) and velocity confidence (right, the coherence of each cell’s velocity with its neighbors) are shown on the UMAP. (**D**) The velocity pseudotime state of each of the three cell types is shown as a violin plot with a black bar indicating the median value. (**E**). The velocity confidence for each cell type is shown; and the dashed line marks the level below which an arrow direction is not interpretable. (**F**). The kernel density of the velocity Pseudotime for each of the four treatment arms are shown. All treatments have a similar Pseudotime distribution, but the SAHA-treated cells have a more differentiated population. (**G**) The mean velocity Pseudotime for each of the 3 cell types are shown for each treatment arm. The dots show the values for each cell type / treatment obtained on each of the two harvest days. (**H-J**) Cytotrace 2 indicates that Mes^Inh^ cells have the lowest developmental potential (less than MyoF or HSC) and are terminally differentiated. (**H**) A bar graph showing the mean CytoTRACE2 potency of each cell type; the 3 cell types of interest are in bold and colored, while other cell types are grey; and the values are shown above each bar. (**I**). The per-cell Cytotrace 2 potency is shown on the reference UMAP. The 3 cell types of interest (8,688 cells) are the dots that are colored according to the scale on the right, and the 51,312 other cell types are shown in grey. The color scale is clipped to show the 2nd–98th percentile of cells (0.064–0.145) for visualization purposes. (**J**). The Cytotrace 2 determined potency distributions of the three cell types are shown for each treatment arm. (**K-M**) Monocle 2 Pseudotime analysis confirms that Mes^Inh^ cells are terminally differentiated. The Monocle 2 determined developmental time progresses from HSC (0.15) to MyoF (1.98) and reaches its peak for Mes^Inh^ cells (3.41). (**K**) The Monocle 2 pseudotime kernel density (median) is shown for the three cell types analyzed. (**L**) The pseudotime axis is shown for each of the four treatment arms. (**M**) The fraction of each cells in a treatment arm that occupies Discriminative Dimensionality Reduction Tree (DDRTree) state 3 is shown. The SAHA-treated arm has the highest value, which indicates that SAHA treated cells are concentrated in the Mes^Inh^-containing region of the tree.

## Discussion

Although HDACi, BRD4i, TGFRB1i, GSK3βi and p38i all reduced fibrosis when co-administered with TGFβ1, a detailed analysis using the microHO system revealed that these drugs had quite distinct pharmacodynamic properties. The BRD4i did not reduce an ongoing fibrosis; p38i and GSK3βi were unable to promote retention of liver metabolic functions, which indicates that reducing fibrosis does not necessarily restore liver metabolic “health”. Only the HDACi and TGFRB1i could reduce an ongoing fibrosis, promote liver regeneration and maintain liver metabolic functions. TGFRB1 inhibitors are not viable drug candidates. Although the cardiovascular toxicity observed in animal models (*48*) could be reduced by dose adjustment, TGFRB1i still causes significant side effects in humans (*49*). Since Vorinostat is an FDA approved drug with the best pharmacodynamic profile of the tested drugs, it is a better candidate for a new liver fibrosis treatment. Moreover, Vorinostat exerts its profound disease-modifying effects at a concentration (1 µM) that is safely and routinely achieved in the plasma of human patients receiving standard oral doses. Oral 400 mg or 800 mg doses of Vorinostat achieved peak plasma concentrations of 1.21 uM (*50, 51*) or 4.8 uM (*52*), respectively, in treated patients. Hence, transient exposures produce Vorinostat concentrations that were several times higher than were anti-fibrotic in microHOs, and these concentrations were clinically tolerated. Vorinostat undergoes heavy first pass metabolism in liver, and patients with liver dysfunction require reduced Vorinostat doses (100 – 300 mg) due to delayed clearance (*53*). Hence, fibrotic liver tissue is uniquely positioned to achieve robust, localized therapeutic concentrations. Although the translational validity is uncertain, HDACi have been shown to decrease injury and/or fibrosis in rodent models of liver (*54–56*) and lung (*57*) fibrosis. Hence, data from animal models that would support the use of HDACi for treating liver fibrosis was already generated.

*Why was Vorinostat uniquely able to reduce an ongoing fibrosis and counter TGFβ1-induced repression of liver metabolic functions?* Our data suggest that the inability of the BRD4i to reduce an established fibrosis—despite matching SAHA’s efficacy when co-administered prophylactically with TGFβ1—reflects the difference between transcriptional interruption and epigenetic reprogramming. In an established fibrotic microenvironment, halting active transcription could be insufficient to reduce the ECM or dismantle established epigenetic silencing. The differential effects of BRD4i and HDACi likely operate across three distinct mechanistic levels (**Fig. 7**).

**Figure 7.**
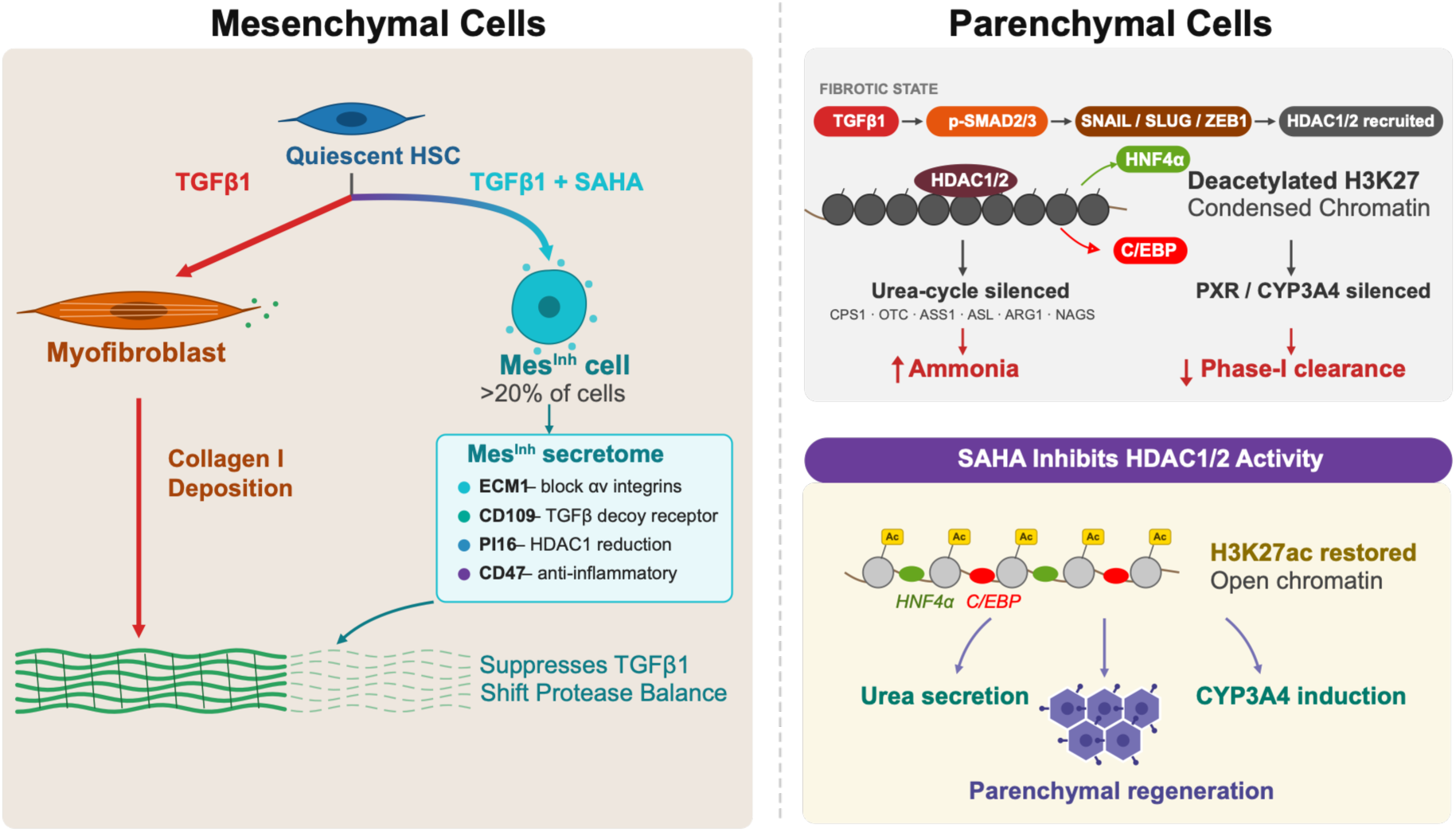
A diagram of the anti-fibrotic effects of Vorinostat in the mesenchymal (Left) and parenchymal (right) liver cell compartments. TGFβ1 induces HSC to become activated myofibroblasts that produce excess ECM. Vorinostat redirects HSC differentiation into Mes^Inh^ cells (∼20% of cells in a microHO) that express multiple, potent anti-fibrotic effectors that can resolve the fibrotic matrix produced by myofibroblasts. In the parenchymal compartment of a fibrotic liver, TGFβ1 signaling recruits HDAC-containing complexes that deacetylate histones, which condense and inactivate chromatin at the sites required for liver specification (acted upon by HNF4a and C/EBP) and for essential liver metabolic functions (ammonia and phase 1 (CYP3A4-mediated) drug metabolism). Vorinostat inhibits HDAC activity, which restores histone acetylation patterns, and opens the chromatin at key genomic sites that are bound by liver fate determining pioneer and other transcription factors. Because of its epigenomic effects, Vorinostat can reduce myofibroblast generation, preserve liver metabolic functions and promote liver parenchymal cell regeneration.

First, SAHA does not simply suppress MyoFs; it actively redirects HSC differentiation into Mes^Inh^ cells that comprise >20% of the cells in a microHO, which we have shown to express potent endogenous anti-fibrotic effectors (*22*). These include: ECM1, a protein that binds αv integrins to maintain TGFβ in an inactive state (*58*); peptidase inhibitor 16 (PI16), which attenuates fibrosis via regulation of fibroblast HDAC1 (*59*); CD47, a protein that inhibits liver fibrosis in mice (*60*); and CD109, a TGFβ coreceptor antagonist that inhibits pulmonary fibrosis in mice (*61*). The Mes^Inh^ secretome could actively neutralize ongoing TGFβ signaling and could alter the local protease/hydrolase balance to promote ECM clearance. Although BRD4i can freeze or blunt ongoing transcription, it does not drive HSC transdifferentiation into matrix-resolving mesenchymal cells. Vorinostat’s ability to do this is consistent with the capacity of anti-cancer drugs to act as cellular differentiating agents (*62–64*); and Vorinostat specifically has been shown to induce mesenchymal cell differentiation (*65*).

Second, in an ongoing fibrosis, TGFβ1 utilizes the SMAD2/3 signaling cascade to actively repress master transcription factors (HNF4α and C/EBP) that function as pioneer factors to maintain open chromatin at the enhancers of liver-specific genes(*66–68*). TGFβ1 also promotes the expression of epithelial-mesenchymal transition (EMT)-inducing transcription factors (*SNAIL, SLUG, ZEB1*) that recruit co-repressor complexes containing HDAC1/2 to silence urea cycle enzymes (*69, 70*) and *PXR/CYP3A4* expression (*71, 72*). By directly blocking HDACs, SAHA can open condensed chromatin, de-repress liver metabolic loci, and restore the EPCAM^+^ parenchymal population, enabling maintenance of ammonia metabolism and Phase-I drug metabolism capabilities. In contrast, BRD4 is a reader that binds to existing N-acetylated lysine residues (such as H3K27ac) (*73, 74*) to facilitate RNA Pol II promoter proximal pause-release (*75, 76*). It is fundamentally unable to de-repress promoters and enhancers that have already been deacetylated and condensed by TGFβ1-recruited HDAC complexes. This is consistent with our finding that Vorinostat co-administration reduced TGFβ1-induced chromatin structural changes in microHOs by 91%; whereas BRD4i co-administration had a lesser effect (43% reduction) (*22*), and the BRD4i effect on chromatin structure would be much smaller if administered during an ongoing fibrosis.

Third, BRD4 cannot reverse the dynamics of an already established fibrotic matrix. When co-administered prophylactically with TGFβ1, a BRD4i can prevent the initial assembly of super-enhancers at pro-fibrotic effector genes (*COL1A1*, *ACTA2*) and can prevent quiescent HSCs from transitioning into active collagen-secreting MyoFs. However, when crosslinked type I collagen bundles have already formed across the organoid lobule, a BRD4i can only prevent additional collagen mRNA synthesis, but it cannot affect degradation of existing matrix or dismantling of established pro-fibrotic autocrine loops. In summary, our data indicate that Vorinostat overcomes these barriers to uniquely resolve an existing fibrosis and to “re-open” liver metabolic functions by actively redirecting mesenchymal cell differentiation and reversing TGFβ1-induced epigenetic silencing.

## Acknowledgements

This work was supported by NIH awards (1R01DC021133 and 1 R24 OD035408) to GP. The content is solely the responsibility of the authors and does not necessarily represent the official views of the NIH. We thank Dr. Robert Lewis for careful reading of this manuscript and providing helpful suggestions throughout this project and Dr. Le Cong for providing the cell lineage barcode constructs.

## Data availability

For the DAISY barcode dataset: the raw sequencing reads (WT and EC libraries), per-sub-library count matrices, cell metadata, gene tables and per-molecule transcript assignment tables are available from GEO under accession number GSE349454. The sample sheet and the recorder and reporter reference sequences are deposited with the same series. For the drug atlas: the raw data are available from GEO (accession numbers GSE228214 and GSE306096). The processed object is available from Zenodo (doi:10.5281/zenodo.14996331). Jupyter notebooks and scripts reproducing the barcode and scRNA-seq analyses and figures from the data, with the software environment specification will be made available upon request.

## Author contributions

Conceptualization: G.P., Y.G. Methodology: Y.G., G.P. Investigation and formal analysis: Y.G, G.P. Funding acquisition and supervision: G.P. Writing—original draft: G.P. Writing—review and editing: Y.G, GP.

## Abbreviations

ECM: extracellular matrix
EPCAM: Epithelial Cell Adhesion Molecule
EMT: epithelial-mesenchymal transition
FCM: flow cytometry
FDR: false discovery rate
HSC: hepatic stellate cells
HDAC: histone deacetylase
MOA: metabolism associated steatohepatitis, mechanism of action
MyoF: myofibroblast

