## Supplemental Methods, figures and tables for "Vorinostat Can Reduce an Ongoing Fibrosis and Preserve Key Liver Metabolic Functions by Selectively Altering Mesenchymal Cell Differentiation"

### Supplemental Information

**Supplemental Note 1**: *Characterization of DAISY cell barcodes in microHOs*. To establish the fidelity and controllability of the lineage recorder, we first validated Cas12a expression and cassette editing by analysis of bulk samples. RT-PCR confirmed that Cas12a expression was strictly dependent on 500 ng/ml doxycycline (Dox) induction in both the reporter COL1A1:Clover line and in parental C3 iPSC lines, and it had negligible background leakage expression in uninduced controls (**Fig. S6A**). We profiled editing dynamics across multiple Dox induction schedules in COL1A1:Clover microHOs that were harvested on Day 20, which were benchmarked against unedited controls (pN256 plasmid and uninduced microHOs) (**Fig. S6B**). Extending the Dox exposure period (3 days, 10 days, and 13 days) resulted in the barcodes progressively accumulating target sequence modifications while maintaining low background editing in uninduced microHOs (Fig. S7B). Analysis of Cas12a editing events revealed that there was a broad distribution of unique modifications at the target sites; different indel alleles comprised a reproducible proportion of the sequencing reads across the different treatments (**Fig. S6C**). Quantification of editing entropy demonstrated that there was high diversity and balanced information density across the DAISY target sites, which confirmed that there was sufficient clonal resolution for single-cell lineage reconstruction (**Fig. S6D**).

Before reconstructing cell phylogenies from the single cell DAISY dataset, we evaluated whether the DAISY recorder retained sufficient clonal diversity and editing information across the differentiation stages, which went from iPSCs (Day 0), hepatoblasts (HB, Day 9), early hepatic organoids (eHO, Day 13), and mature hepatic organoids (HO, Days 20–30). After molecule-level filtering, 56,713 cells were recovered, which captured 402,306 UMIs across 9,344 unique integration barcodes (intBCs). The cells had a median of three intBCs (mean 3.68; 13.8% carried a single barcode); whereas 61.2% of intBCs were restricted to single cells, and the top 20 most frequent intBCs comprised 45.8% of cell–intBC pairings (**Fig. S7**). Although the raw intBC richness decreased 4.8-fold between the iPSC and eHO stages, this decline reflected sequencing depth; after rarefying to 22,662 UMIs per stage, intBC richness varied by only 1.22-fold (**Fig. S8**). Furthermore, the overall clonal architecture remained stable between the differentiation stages, with HB and mature HO yielding comparable clone counts (785 vs. 778 clones) and top-eight clonal shares (32.4% vs. 33.4%)(**Fig. S10**).

Of importance, lineage recording progressively accumulated as differentiation advanced. Across the individual target sites, the fraction of edited sites expanded from 1.3–2.5% in iPSCs to 9.3–11.8% in mature HOs (**Fig. S11**), with concordant kinetic increases across all four target sites (**Fig. S9, S11-12**). Rarefied per-site Shannon entropy similarly increased from 0.29–0.47 to 1.15–1.54 bits. Target editing was consistently observed across all mature HO treatment arms (5.5–16.3% per site) (**Fig. S13**), while the editing efficiency varied 18-fold across the 15 most frequent intBCs (0.36–6.52%). The differences between individual clones or treatment arms were not ascribed biological significance due to uneven batch pooling across conditions (with the collection day shifting the HO edited fraction between 7.1% and 22.3%). Thus, barcode diversity is preserved throughout long-term organoid culture without technical bottlenecks; which ensures that there is a robust, depth-controlled barcode recording substrate for phylogenetic reconstruction.

**Methods**

***AI usage disclaimer:*** scRNA-seq analysis including transcriptome, phylogeny and trajectory code was written and executed with the assistance of Claude (Anthropic) that was run through the Claude Code command-line interface. All analyses in Jupyter notebooks were generated programmatically from versioned Python builder scripts that put out the R/Seurat analyses, the statistical tests and the plots, which enable every result to be reproduced by regenerating and re-executing the notebook from its builder. The study's scope, analytical decisions and interpretations were directed by the authors, who verified all the reported numbers against the notebooks' own output tables, and they take full responsibility for the content.

***iPSC line and COL1A1 reporter.*** The iPSCs (C3 and C3 COL1A1-P2A-Clover) used in this paper were maintained feeder-free in StemMACS™ iPS-Brew (Miltenyi Biotec, San Jose CA) on Matrigel and passaged with 0.5 mM EDTA at 1:10 ratio. The human iPSC line (C3 COL1A1-P2A-Clover) used in this study was prepared as previously described (*1*). It has a knock-in P2A-linked fluorescent reporter (Clover) at the 3′ end of the endogenous *COL1A1* locus; and the self-cleaving P2A peptide labels COL1A1-expressing cells with an intracellular fluorescent protein that does not disrupt collagen production. The reporter transcript is represented in the custom alignment reference as the 769 bp contig COL1A1-P2A-HOT and is quantified per cell along with the endogenous transcriptome.

***Hepatic organoid differentiation in microwells.*** To produce microHOs, 50~60% confluent iPSC cells were cultured in a series of differentiation media as previously described (*1, 2*). In brief, reporter iPSCs were differentiated to hepatoblasts (HB) over 9 days. On Day 9, 10,000 hepatoblasts were seeded into a microwell and allowed to aggregate and then differentiate into multi-lineage human hepatic organoids in microwells (microHOs). The cultures were maintained in organoid medium with the medium exchanges performed every 3~4 days; and differentiation continued to the hepatic organoid (HO) stage. These microHOs have hepatocytes, cholangiocytes, bile ducts and lobule-like architecture; and after TGFβ1 exposure they have myofibroblasts and produce collagen-rich thick fibrils (*1, 2*).

***TGFβ1 and drug treatments.*** *Co-administration protocol*: Treatment with TGFβ or TGFβ + drug was initiated on Day 13, and the microHO cultures were differentiated through 21 or 31 days as described (*2*). *Fibrosis reversal protocol:* microHOs were matured until Day 21 before TGFβ1 (50 ng/ml) was added. After one week of fibrosis induction, TGFβ1 and drug were administered to the fibrotic microHOs on day 28, and the cultures were maintained through Day 31 to 35. Experimental conditions included a DMSO vehicle control (NC) and a TGFβ1 arm (50 ng $mL^{-1}$; T) against which drug effects were evaluated. The co-treatments tested consisted of a positive control TGFBR1 inhibitor (A83-01; 10 uM; Ti), a GSK3β inhibitor CHIR-99021 (3 μM; CHIR), a p38 MAPK inhibitor (SB202190 10 μM; P38i), an HDAC inhibitor (Vorinostat or SAHA 1 μM); and a BRD4 inhibitor (ZEN-3694; 2 μM). All drugs were solubilized in DMSO.

***Histological Staining of microHOs***. To evaluate ECM deposition and bile acid production in microHOs from the different treatment groups, the microHOs were processed for specialized staining using commercial assay kits according to the manufacturers' instructions. Collagen deposition and connective tissue distribution were assessed using Masson's 2000 Trichrome kit (American MasterTech, KTMTR2 Lodi CA). Bile pigment detection was performed using the Bile Stain Kit, Halls (American MasterTech, KTHBI).

***Measurement of CYP3A4 activity***. The effect of fibrosis and drugs on CYP3A enzyme activity in microHOs was measured using the P450-Glo™ CYP3A4 Assay (Promega, Madison WI). This assay uses a derivatized luciferin (Luciferin IPA) that is converted by CYP3A4 enzyme activity to a substrate that reacts with luciferase and produces light that is directly proportional to the level of CYP3A4 activity. microHOs were prepared as described in (*1, 2*). (prevention protocol), and TGFβ (50 ng/ml) was added to some microHOs cultures on day 13. On day 21, 100 ul of fresh media containing each experimental drug with 3 uM Luciferin-IPA along with 25 μM rifampicin (which binds the pregnane X receptor and acts as a CYP3A4 inducer); or 1 μM ketoconazole (a CYP3A4 inhibitor) was added to selected microwells. Cultures were incubated at 37°C for 6 or 24 hours, and then 25 ul of the media from each well was transferred to another plate where CYP3A4 activity was measured according to the manufacturer's instructions. Following a 20-minute incubation with the luciferin detection reagent, luminescence was recorded with a 1-second integration time per well using a BMG FLUOstar microplate reader. Each measurement is the average of data obtained from 8~16 microHO that were prepared in 4 experimental batches. Differences between the groups was assessed using the Student's t-test.

***Measurement of urea cycle activity***. microHO cultures were prepared by as described in (*1, 2*). (prevention protocol). Supernatants (50 ul) were harvested from control, TGFβ-treated (fibrotic), or TGFβ + drug-treated microHO cultures on day 21. The supernatants were centrifuged (2,000 x g, 5 minutes) to remove debris and then stored at -80 °C. Urea concentrations were determined using the non-enzymatic Jung method (*3*) (Bioassay Systems, Hayward CA, DIUR-100) according to the manufacturer's instructions. This assay relies on the reaction of urea with diacetyl monoxime under acidic conditions to form a stable, yellowish-orange diazine derivative whose absorbance is measured at 430 nm by a microplate reader. Net urea production was calculated by subtracting the absorbance of a media-only blank, which accounts for baseline interference from other nitrogenous compounds in the supernatant. Each measurement is the average of data obtained from 16 microHOs. The statistical significance between groups was assessed using the Student's t-test compared to TGFβ group.

***DAISY lineage recorder construct and transduction.*** AAVS1 sit Knock-in iCas12a (pQ1_AAVS1_Tet_Cas12_KI, provided by Prof. Cong's lab) (*4*) and lentiviral iCas12a (Addgene: 188471, pN251) iPSC lines were established. The level of doxycycline-induced expression (DOX 500 ng/ml) was confirmed (Fig. S6). The DAISY recorder (pN256, Addgene: 188472) is a single expressed cassette carrying four Cas target sites and a 10 bp random integration barcode (intBC). The 175 bp recorder region is the target reference used for allele calling; and the editing sites lie at positions 45, 61, 105 and 121, and the intBC occupies the degenerate 10-mer at positions 143–153. The cassette was delivered to iPSCs by lentiviral transduction before Day 0 at a multiplicity of infection (MOI) of 3, which was selected to produce a small number of independent integrations per cell. The actual number of intBC in the sequenced libraries had a median of 3 per cell (mean 3.68). Because delivery precedes Day 0, no integration is acquired later in the experiment, and every intBC observed at the mature organoid stage was present at Day 0. Transduced cells were expanded before differentiation. Four independent transductions were performed, yielding the four batches analyzed here. Since each batch was barcoded separately, no clone is shared between batches, and all clonal analyses were performed within a batch. For a single cell (sc)DAISY experiment, the co-administration protocol was used, and Cas12a expression was induced immediately after day 9 in HB cultures by addition of 500 ng/ml Dox. Four independently transduced and independently barcoded batches (Batch 1, 2, 4 and 6) were sampled across four differentiation stages: iPSC (Day 0), HB (Day 9), early HO (eHO, Day 13) and mature HO (Days 20, 21, 26 and 30). The iPSC, HB and eHO samples precede or coincide with the start of a treatment, and they therefore represent untreated baselines. Only the mature organoid samples carry a treatment arm. Cells were fixed and permeabilized according to the Parse Biosciences Evercode Cell Fixation protocol (Seattle, WA) and were stored at −80 °C prior to library preparation.

***Single-cell library preparation and sequencing.*** Libraries were prepared using the Parse Biosciences Evercode WT split-pool combinatorial barcoding kit (CRISPR Detect for Evercode™ WT v3, Seattle WA) according to the manufacturer's instructions. Each sample was loaded into its own reverse-transcription well of the round-1 barcoding plate (i.e., the round-1 barcode identifies the sample); and cells then underwent two further rounds of split-pool ligation barcoding. The barcoded material was divided into eight sub-libraries, and the mapping of each sample from the reverse transcription well to that batch, treatment, day and stage is recorded in meta_DAISY.csv. The recorder is expressed and captured in the same cDNA, but it is present at a level that is too low for confident allele calling from the whole-transcriptome library. Therefore, a targeted amplicon library was separately prepared from the same barcoded cDNA using cassette-specific nested forward primers (P1: *CTTCTGCTACGTCCCTTCGG*; P2: *TCGTCGGCAGCGTCAGATGTGTATAAGAGACAGCCATCTCATCCCTGCGTGTCTCC*) that were paired with the Parse adapter; and this preparation was used with 19 cycles of PCR followed by bead clean-up. Each sub-library generated two sequencing libraries from the same cells: a whole-transcriptome (WT) library for single-cell RNA sequencing and a lineage-barcode enrichment (EC) library for barcode analysis. Since the DAISY primer mis-primes at a detectable rate, a substantial fraction of the resulting library can align to off-target sites. The mis-primes are removed computationally using the flank screen and alignment-score filter described below. Amplicon libraries were paired end sequenced with a 150 bp read 1. Since this reaches position 150 of the cassette and stops three bases short of the end of the integration barcode; merged read pairs were extended to 227 bp. That read geometry is the basis for the truncated-barcode class handled in preprocessing.

***Hepatic organoid drug atlas (reference).*** The reference atlas is a scRNA-seq atlas of microHOs treated with TGF-β1 alone or combined with anti-fibrotic compounds (*1, 2*). It holds 613,826 cells from 17 experimental batches, which contains 16 annotated cell types (*1, 2*). The raw data is in: GEO GSE228214 and GSE306096. The processed object derived from them (*2*) was deposited at Zenodo (doi:10.5281/zenodo.14996331; file scRNA-seq_all_drugs_noB1921_processed.rds). The 16 cell-type labels were renamed as: eEC, EC, Progenitor, preCho, Cho, Hep1, Hep2, Hep_WNT, EMT, HSC, MyoF, $Mes^{Inh}$, Mes_WNT, Mes1, Mes2 and Mesothelia (table atlas_celltype_recode.csv).

***Whole-transcriptome libraries.*** A custom human genome reference was built with split-pipe --mode mkref (Parse Biosciences split-pipe v1.7.3) from: - the GRCh38 primary genome assembly with Ensembl release 113 annotation; and the 769-bp *COL1A1-P2A-HOT* reporter sequence was added as an additional contig and gene. Each WT sub-library was processed with split-pipe --mode all --kit WT --chemistry v3. This demultiplexes the three-round cell barcodes and UMIs, aligns cDNA reads with STAR 2.7.10a , and counts unique transcripts per gene and cell. For every sub-library we used: the filtered cell-by-gene count matrix (DGE_filtered) and the per-molecule transcript assignment table (tscp_assignment.csv.gz). Four arms of the atlas were analyzed: vehicle (NC), TGF-β1 (T), TGF-β1 + SB431542 (Ti) and TGF-β1 + vorinostat (SAHA). A well is one batch × arm (one sample per well). Each drug was evaluated in batches that contained NC, T and T+drug: SAHA was used in seven batches (B47, B50, B52, B54, B55, B56, B57); and Ti was used in seven batches (B14, B26, B30, B50, B51, B52, B55). For each well and cell type, the share of each cell type in the well was calculated, and the was transformed to a $\log_2$ odds with a Haldane correction of 0.5 (to eliminate the ‘Zero Cell’ problem).

***The rules for two contrast comparisons.*** For each cell type and experimental batch, each cell type’s proportion in the well of that batch was calculated for each treatment arm and converted to $\log_2$ odds. Two contrasts were formed *within* each batch: contrast 1: TGFβ versus vehicle (T − NC); and contrast 2: TGFβ versus TGFβ plus drug (T − drug). Hence, each batch contributed one paired value per contrast and the between-batch variation in baseline composition was cancelled. The significance was assessed on samples rather than on cells, using a paired two-sided $t$-test on the seven per-batch differences for each contrast and cell type. The two contrast comparisons were tested in sequence: contrast 1 was corrected across all sixteen of the identified cell types using the Benjamini–Hochberg procedure, and contrast 2 was then tested only within the cell types where contrast 1 reached a FDR ≤ 0.05 and were corrected across that smaller set because a drug effect is interpreted relative to a TGFβ effect and is undefined where there is none. A cell type was flagged when both contrasts were non-zero, with no constraint on the direction. A tier A result required FDR ≤ 0.05 for both contrasts, while a tier B result had a FDR ≤ 0.05 on contrast 1 with nominal $p \leq 0.05$ on contrast 2. The direction of change in cell number was recorded; a cell type whose two contrast estimates share a sign was labelled *reverted* (i.e., the drug returns it toward that in the vehicle) and one whose estimates differ in sign *amplified* (the drug moves it further than did TGFβ treatment alone). A Haldane correction of 0.5 was added to every cell count before the log-odds were calculated so that an experimental arm containing none of a cell type remained finite, and the fraction of the TGFβ shift reversed, rev = $(p_T - p_{drug})/(p_T - p_{NC})$, was reported descriptively and entered as a no call.

***Reference mapping of DAISY organoid cells onto the atlas.*** The Batch 6 microHO cells (seven arms, Days 21 and 26; 81,863 cells) of the DAISY dataset were mapped onto the drug atlas using Seurat 5.5.0 (*5*). The atlas's 2,000 variable features that were present in the query (1,751 genes). FindTransferAnchors with normalization.method = "LogNormalize", reference.reduction = "RPCAIntegration" and dimensions 1–30. Defaults applied: k.anchor = 5, k.score = 30, k.filter = NA. - Transfer: MapQuery transferred the 16 cell-type labels. Each query cell received the label with the highest prediction score, plus its score. Cells were projected into the atlas UMAP through the re-fitted model. The per-arm composition of the query was computed from the transferred labels, will data pooling from both harvest days.

***Lineage-Barcode Processing and Allele Calling***. For read processing: raw read pairs were parsed to extract UMIs and Evercode cell barcodes from read 2, which was reverse-complemented and merged with read 1 based on sequence overlap (minimum 85% identity). Merged reads were written as *unmapped BAM* records with the raw cell barcode, UMI and sub-library read group. Barcodes were corrected against the Evercode v3 whitelist (Levenshtein distance ≤ 1), matched to validated transcriptomic barcodes, and formatted for single-cell matching. Allele calling was performed using Cassiopeia (*6*); the reads were collapsed by cell barcode and UMI, resolved to consensus sequences, and filtered to exclude multiplets (>1,000

UMIs or >100,000 reads). The sequences were aligned to the 274-nt recorder reference amplicon using the Smith–Waterman alignment (score ≥ 200). Indels were called within ± 8 nt of Cas12a target sites alongside the 10-nt integration barcode (intBC). Following homopolymer removal, UMI error correction, and doublet removal (>35% discordant alleles), each cell–intBC pair was resolved to a single dominant allele. For Lineage Tree Reconstruction and Mesenchymal Dynamics Phylogenetic Reconstruction, separate trees were built for Day 21 and Day 26 samples from Batch 6 cells across the four treatment arms (NC, T, Ti, and SAHA). Cells were clustered into lineage groups by shared intBC sets using optimized thresholds. Character matrices (intBC x cut site) were solved per clone with Cassiopeia's VanillaGreedySolver, and clone trees were joined under a synthetic root. For mesenchymal lineage analysis (HSC, MyoF, and $Mes^{Inh}$), the inter-type distances were quantified as the mean number of edges separating leaves through their lowest common ancestor. To control for leaf depth and treatment arm confounders, observed tree distances were evaluated against a stratified permutation null model (500 permutations) and hierarchically clustered by average linkage with bootstrap resampling (200 replicates). Evolutionary coupling was computed using Cassiopeia's permutation model and benchmarked against the depth- and arm-controlled metrics.

***RNA velocity, differentiation potency and pseudotime analyses.*** For RNA Velocity, the spliced and un-spliced reads were quantified from molecule tables. Using scVelo (*7*), the velocities were estimated for the HSC, MyoF, and $Mes^{Inh}$ subsets by computing moments across 30 nearest neighbors in 30 principal components under the stochastic model. Velocities were projected as streamlines on the UMAP, evaluated as mean grid vectors (22 × 22) for NC, T, Ti, and SAHA arms; and used to derive velocity confidence and pseudotime distributions across cell states and treatment arms.

For differentiation potency: CytoTRACE 2 (*8*) was applied to raw counts from 20,005 cells sampled across cell types and treatment arms and filtered for genes that were detected in at least 10 cells. The relative differentiation potency scores were computed and projected onto the reference UMAP to determine the developmental hierarchy. Monocle 2 Pseudotime Trajectory inference (*9*) was performed on 4,992 CytoTRACE 2-scored cells across 827 shared variable genes using a negative binomial model with DDRTree dimension reduction. The tree was rooted in the HSC-enriched state possessing the highest mean differentiation potency. Pseudotime trajectories and state densities were evaluated across cell types and treatment conditions, which specifically assessed branch occupancy in the $Mes^{Inh}$ enriched state.

### Supplementary figures

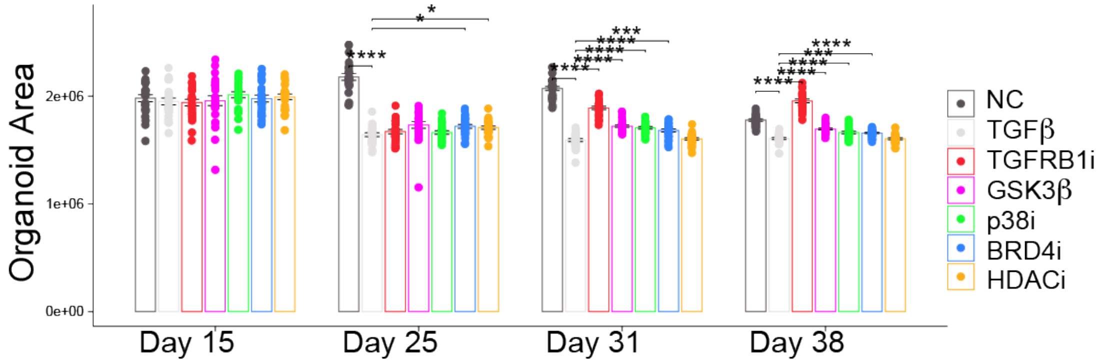


**Figure S1**. microHO size was serially measured on days 15 through 38 in microHOs treated with TGFβ1 or with TGFβ1 + the indicated drugs (as in figure 1) to assess treatment toxicity. TGFβ1 induced a significant decrease in microHO size relative to normal control (NC) microHOs on days 25, 31 and 38. In contrast, all the tested drugs didn't induce further microHO toxicity. Statistical significance was assessed using a student's t-test * $p < 0.05$, *** $p < 0.001$, **** $p < 0.0001$. Control and all other treatment groups were compared with TGFβ1-treated microHOs.

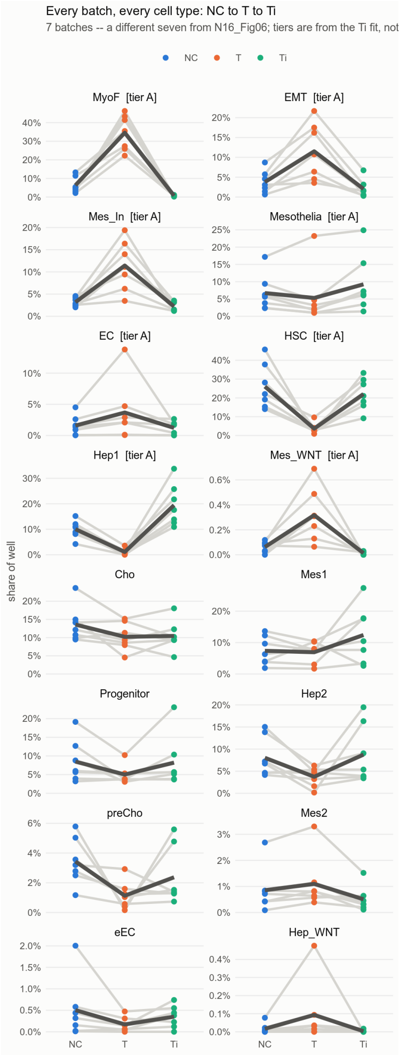


**Figure S2.** Graphs comparing the changes in the percentage of all 16 cell types in microHOs after treatment with control (NC), TGFβ1 (T), or TGFβ1 + TGFBR1i (Ti). It shows the data obtained from every experimental batch and cell type for the two contrast **NC** → **T** → **Ti** comparison. Each group uses microHOs prepared from 7 separate experimental batches and a Tier A label indicates significant differences were noted in both contrasts. There is one grey line per batch, the dark line is the mean across the seven batches. The *y* axes are free, so the different panels are not comparable to each other in height. The data shown in Figs. 4C-D was obtained from these analyses.

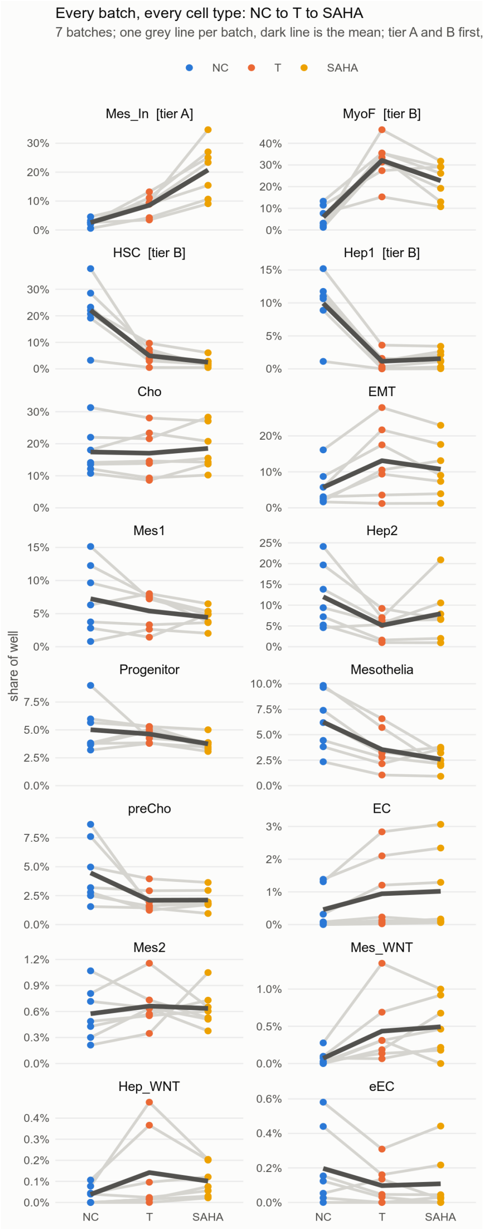


**Figure S3.** Graphs comparing the changes in the percentage of all 16 cell types in microHOs after treatment with control (NC), TGFβ1 (T), or TGFβ1 + SAHA. It shows the data obtained from every experimental batch and cell type for the two contrast **NC → T → SAHA** comparison. Each group uses microHOs prepared from 7 separate experimental batches. Tier A result requires that the FDR ≤ 0.05 for both contrasts, while a Tier B result had a FDR ≤ 0.05 for contrast 1 with nominal $p \leq 0.05$ for contrast 2. There is one grey line per batch, the dark line is the mean across the seven batches. The *y* axes have different values, so the different panels are not comparable to each other in height. The data shown in Figs. 4C-D was obtained from these analyses.

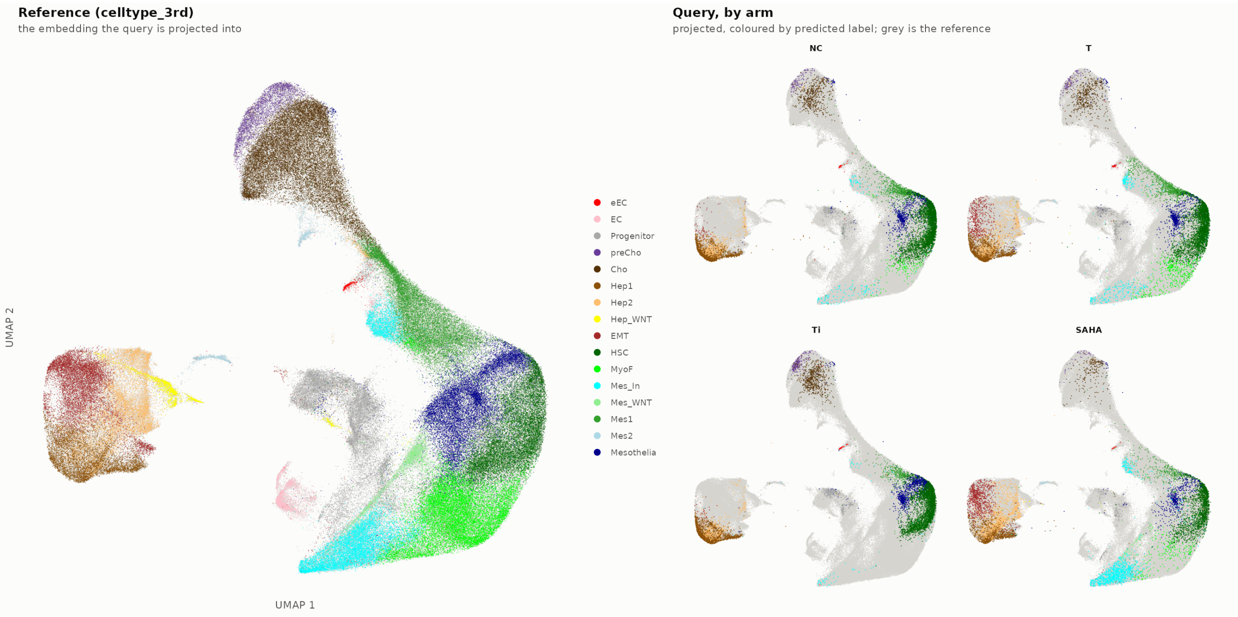


**Figure S4.** microHO cells with a DAISY barcode are projected onto the reference atlas to show the effect of TGFβ1 and coadministration of Ti or SAHA on mesenchymal cells**.** *Left:* The 613,826 microHO cells form the reference atlas, which was generated from a fourteen-arm fibrosis drug-treatment panel, is drawn on its integrated RPCA UMAP and the sixteen curated cell-types are color coded as indicated. *Right:* The 81,863 cells with a DAISY barcode are projected onto the same reference panel embedding with one panel per treatment arm (NC vehicle, T, TGFβ1; or TGFβ1 with Ti (SB431542) or SAHA). The color labeled cell are placed over a grey background of all reference cells. The treatment arms are reduced to 12,000 cells each so that no arm dominates by cell number; and all seven arms appear in the full composition table. Every panel is drawn at a 1:1 data aspect, so distances are comparable between the two axes. These panels clearly show that the bright green MyoFs are present in NC, expand in TGFβ1 microHOs, are essentially absent in Ti, and are present at a reduced number with SAHA co-administration. In contrast, the cyan colored $Mes^{Inh}$ cells are markedly expanded when SAHA is co-administered.

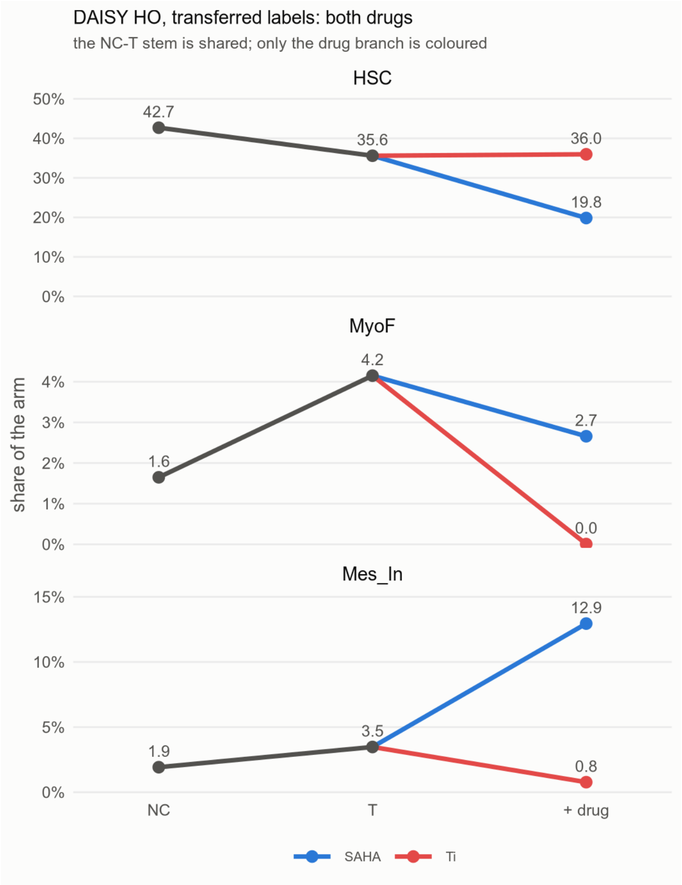


**Figure S5.** Graphs comparing the changes in the percentage of three mesenchymal cell types (HSC, Mes$^{Inh}$, MyoF) in microHOs after treatment with control (NC), TGFβ1 (T), or TGFβ1 + drug (SAHA or Ti). The scDAISY microHO data was used for this analysis. The labels for the treatment groups are: NC → T → SAHA (blue) and NC → T → Ti (red). Since the NC-> T are the same in both series, the stem of the line is black, and the drug treatment branch is colored. The y-axes have different values, so the height of the different panels cannot be compared.

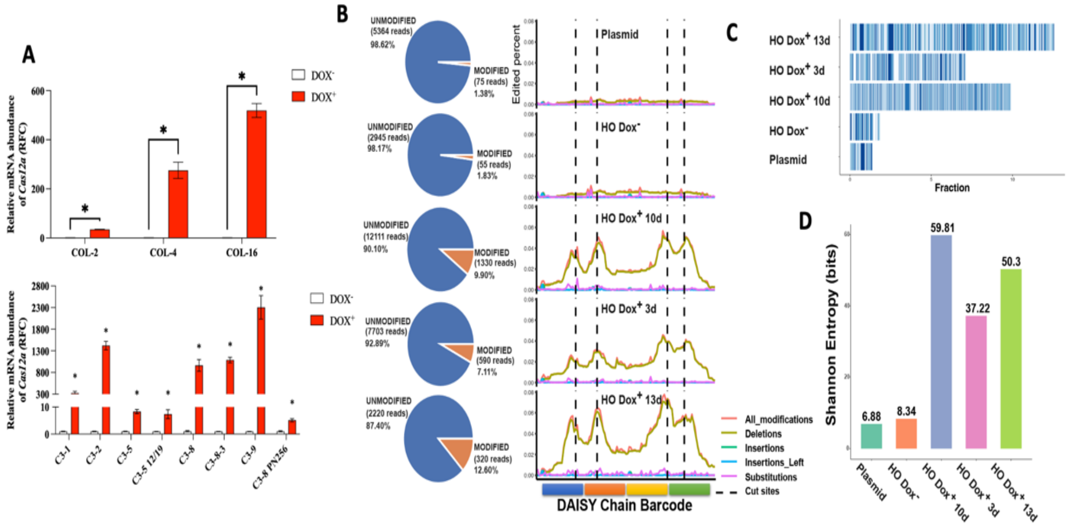


**Figure S6.** Validation of DAISY barcode cassette integration and target-site editing through analysis of bulk samples. (**A**) RT-PCR data showing that Cas12a expression is tightly controlled by Dox treatment in both COL1A1:Clover and C3 cell iPSC lines. The cells were induced by incubation with 500 ng/ml Dox for 2 days. The level of expression of *Cas12a* in control and Dox-treated COL1A1::Clover cells (Top) and in control and Dox treated parental C3 cells (Bottom). (**B**) DAISY editing results in COL1A1:Clover microHOs treated with the negative control (pN256) plasmid; and microHOs without Dox Treatment on Day 20 (HO Dox⁻). The results from microHOs treated with Dox from day 7 to 10 (3d), day 10 to day 20 (10d) or days 7 to 10 (13d) are shown. All microHOs were collected on day 20. The x-axis shows the 4 editing sites in the Daisy barcode, and the y-axis shows percent of edited reads. (**C**) A Stacked bar chart comparing the Cas12a editing outcomes for the barcodes in the microHOs listed in (B). The bar size corresponds with the percentage of bar codes (sequence reads) with a unique modification. (**D**) The calculated Shannon Entropy for Cas12a editing outcomes at DAISY targets in the microHOs described in B.

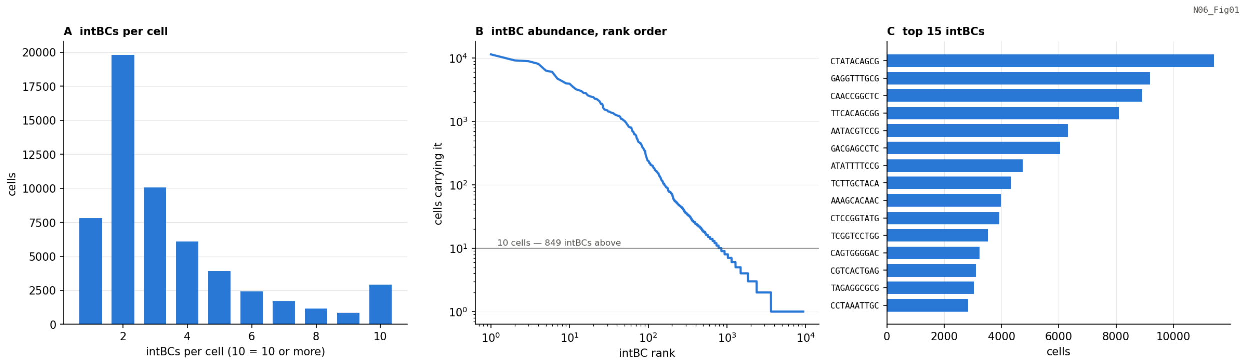


**Figure S7.** The distribution of the DAISY integration barcodes (intBC). The reads from the complete barcode table for the DAISY barcode single-cell experiment had 402,306 barcode-bearing UMIs, 56,713 cells, 9,344 distinct 10 bp intBC, and every read is a full-length construction. (**A**) The distribution of the number of intBC per cell, and the last bar is a pooling of cells with 10 or more intBC. The median is 3 and the mean 3.68 (maximum 116). Note: 13.8% of cells carry one barcode and therefore does not have a clone signature of its own because clone calling matches the co-occurrence of barcodes*.* Hence, a single-tagged cell does not have co-occurrence pattern to match. Also, 22.9% of cells carry five or more intBC. (**B**) The rank order of every barcode based upon the number of cells carrying it. The is a log-log graph, with the 10-cell line marked. (**C**) The number of cells with the fifteen most widely carried barcodes; the top twenty barcodes cover 45.8% of all cellular barcode pairs.

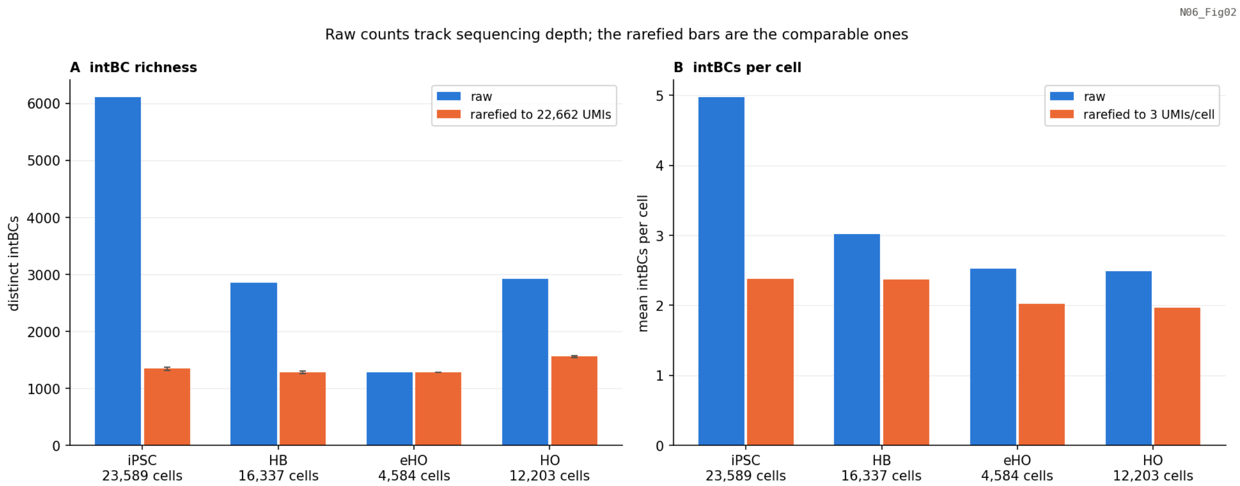


**Figure S8.** Clonal barcode diversity does not decline with microHO differentiation**.** (**A**) The number of distinct intBC per cell (**A**) and the mean number of barcodes per cell (**B**) are shown for each differentiation stage (iPSC, HB, eHO, HO). In both panels, the raw bar is plotted beside a rarefied bar. In panel A, the rarefaction draws the same number of UMIs from every stage before distinct barcodes are counted; and in panel B it draws the same number of UMIs *per cell*, which enables cells with unequal sequencing depth to be compared. Only the rarefied bar is comparable across the different differentiation stages. The stages differ ~10-fold in depth (iPSC has 240,883 UMIs over 23,589 cells; and eHO 22,662 over 4,584), which is why the raw bar richness falls 4.8-fold from iPSC to eHO. In contrast, the rarefied richness differs by only 1.22-fold across all four stages (1,281–1,559 barcodes). The same consideration holds for intBC per cell, which tracks UMIs per cell almost exactly (raw 4.97 → 2.49; rarefied 2.38 → 1.97).

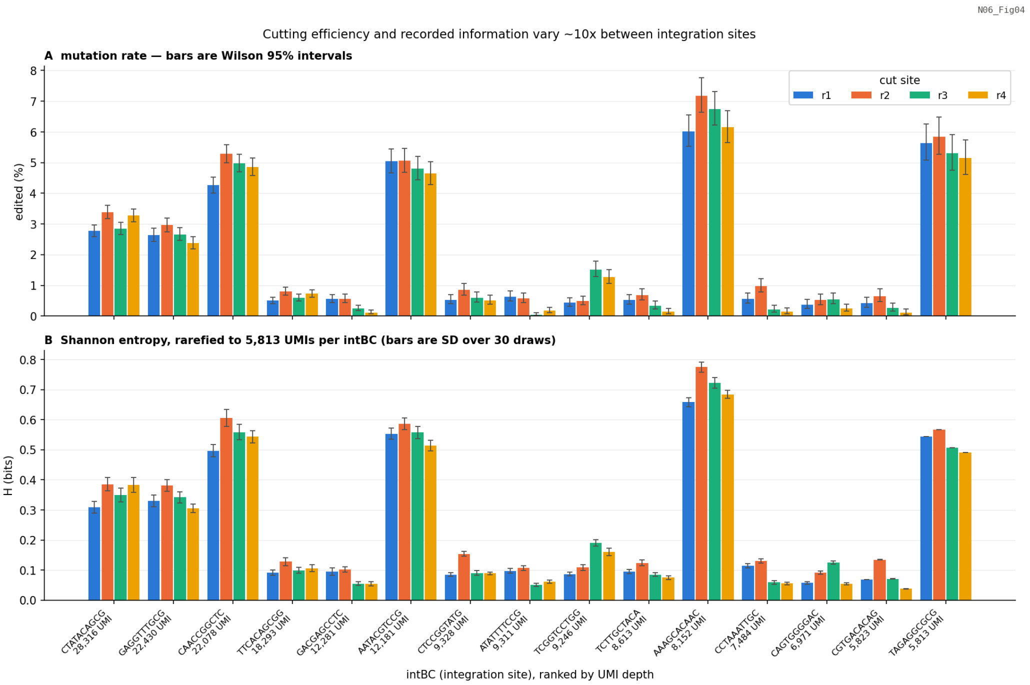


**Figure S9.** The effect of integration position: identical barcode cassettes are edited at ~14-fold different rates depending on where they integrated**.** For the fifteen highest-depth intBC, the fraction of that barcode's UMIs edited at each of the four 4 target cut sites (r1–r4) (**A**), and the depth-rarefied Shannon entropy of that site's allele states are shown in bits (**B**). The edit rate has a ~14-fold range: from 0.36% of UMIs (`GACGAGCCTC`) to 4.89% (`AATACGTCCG`). The rarefied per-site entropy runs 0.04–0.78 bits against a fair coin's 1.0 bit. The four target sites within a barcode behave similarly to one another (site r2 is the most edited in most barcodes), so the variation is caused by different sites of barcode integration rather than between positions in the barcode array.

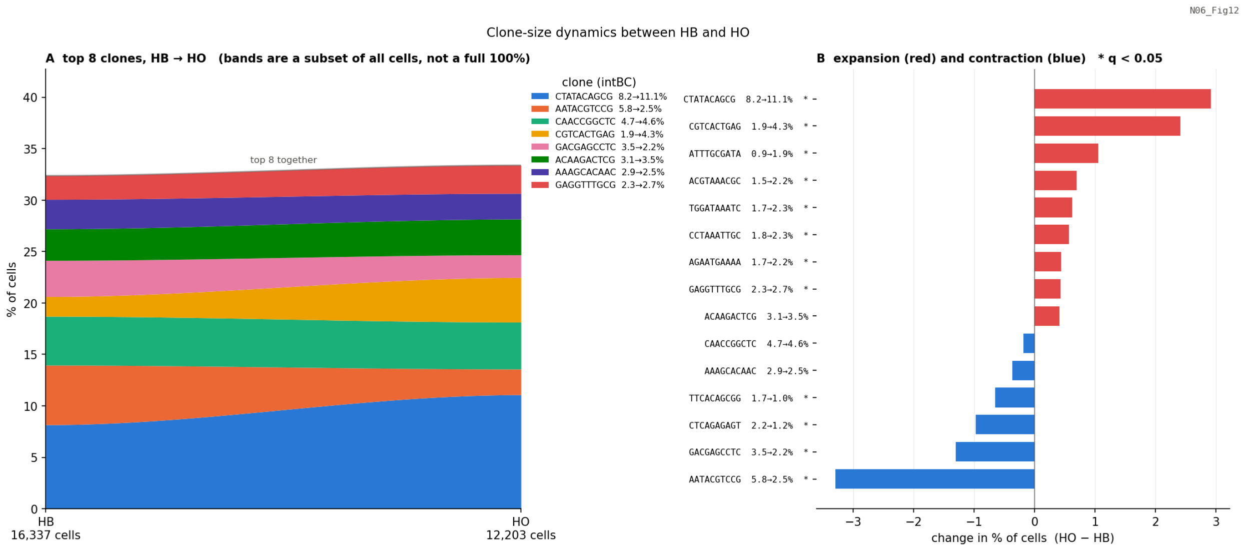


**Figure S10.** While the number of cells in a clone varies between the hepatoblast and organoid stage, the cellular composition is not dominated by any one clone**.** Clone size dynamics from HB (Day 9) to mature microHO (Days 20–30) are shown, which uses pooled data obtained across experimental batches. Here, a clone is defined by its integration barcode. Since the barcode cassette is delivered once before Day 0, the cells sharing a barcode are clonally related. Because a cell carries a median of 3 barcodes, the per-barcode proportions would sum to far more than 100% and could not be stacked together. Hence, each cell is assigned to a single clone, which is its highest-UMI barcode, to make the proportions sum to 100% per stage and every cell is counted once. (**A**) A Fish (Muller) plot of the top clones. The top eight clones cover 32.4% of HBs and 33.4% of mature microHO cells, and the remaining ~1,250 clones account for the other two-thirds of the cells. (**B**) The same clones are shown as expanding (red) or contracting (blue) between two differentiation stages. A star indicates that the comparison has a $q < 0.05$ from a Fisher exact test performed on the cell counts with Benjamini–Hochberg correction. The percentages for twelve of the 15 top clones change significantly between the hepatoblast and microHO states: 8 clones expand and 4 contract. The largest changes are for the `CGTCACTGAG` (1.92 % → 4.33 %, OR 2.32) and `AATACGTCCG` (5.80 % → 2.50 %, OR 0.42) clones. The total clone count is essentially unchanged (785 → 778).

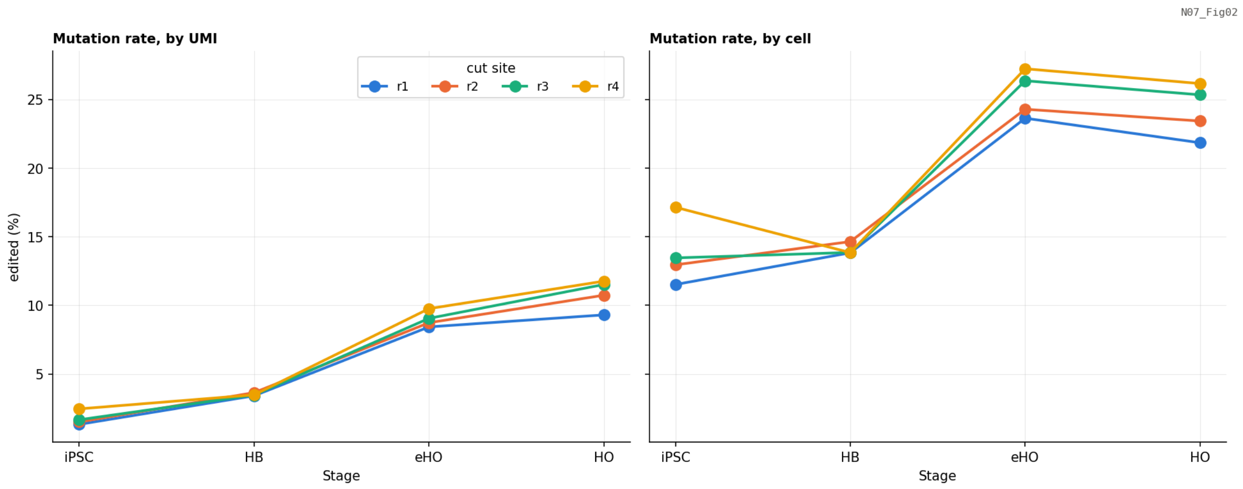


**Figure S11.** Recorder editing at all four target sites in the DAISY barcode increase monotonically throughout microHO differentiation**.** The fraction of edited molecules at each target site (r1-r4) in the DAISY barcode at the iPSC, HB, eHO and mature microHO stages were determined and normalized per UMI (left) or per cell (right). Editing rises at every site (irrespective of the normalization denominator): there is a roughly six-fold increase in the edited fraction from Day 0 to the organoid stage, with site r4 consistently being the most edited and r1 the least. When editing is averaged over the four sites, the means at each stage are 1.75%, iPSC, 3.51% HB; 9.00% eHO, and 10.83% mature microHO. The increase in editing is a property of the barcode since it is maintained at every depth control. However, editing in this system is not cumulative: the recorder fires once and destructively at each site, and only about 1% of later alleles extend an earlier edit, so the edit load is a record of how long Cas12a has had access to the barcode, but it is not a clock that can be read as elapsed divisions. One artefact that is quantified rather than hidden: 22.2% of iPSC edited UMIs sit in a poly-A context that is consistent with oligo-dT internal priming, which is reduced to 1.9–3.7% at the later stages. Since the rate of iPSC editing is likely to be an overestimate, the actual increase in editing with differentiation is likely larger than shown here.

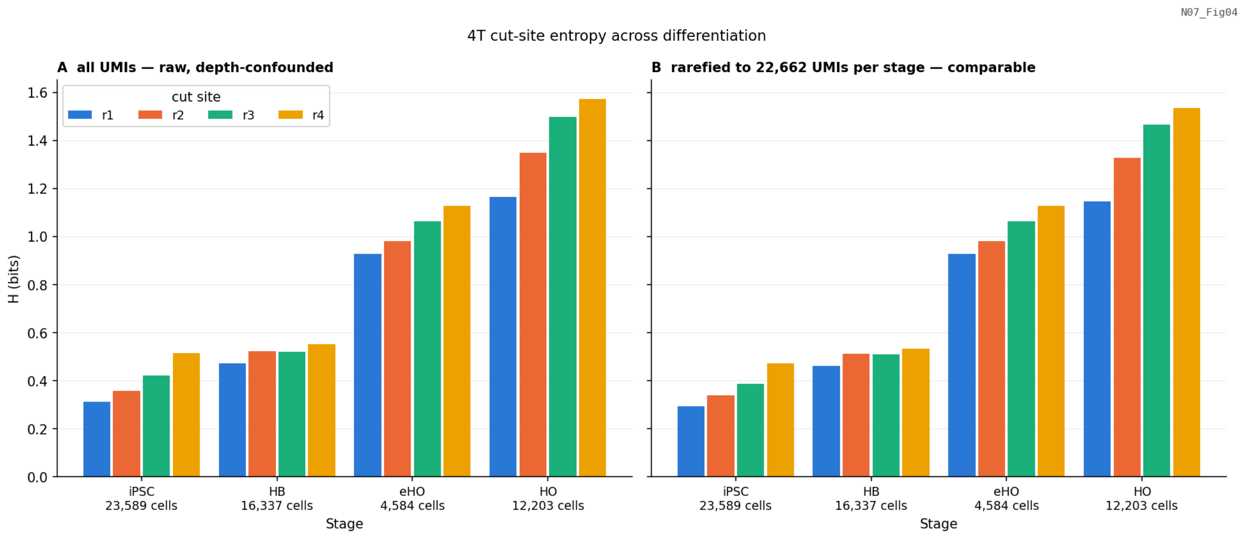


**Figure S12.** The lineage information in a barcode grows 3.7-fold as the microHO differentiates and remains well under one bit per site. The Shannon entropy ($H = -\Sigma\, p_i \log_2 p_i$) over the allele states at each of the four cut sites is shown in bits with UMI-weighted state frequencies; a fair coin is 1.0 bit and a site with one state is 0. **(A)** Shows all states, which is what a tree solver sees (i.e. the realized information). **(B)** The same measure is rarefied to a common UMI depth across the differentiation stages. Since entropy increases with sampling depth ($H \leq \log_2$ of states observed) and the stages differ by ~10-fold in depth, the right panel is more informative. Rarefied entropy rises ~3.7-fold from the iPSC stage (0.29–0.47) to the mature microHO (1.15–1.54 bits per site). The absolute level is the important number for interpreting each tree. Even at the mature HO stage a cut site carries about 1.4 bits, because the unedited state is most common at every site, and the mean over all barcodes of the whole-state entropy is only 0.26 bits. When the unedited state is excluded, the entropy of the indel spectrum *for* a cut is 4.6–7.4 bits (depending upon the stage), which corresponds to roughly 11.6 effective indel states per cut.  Hence, the cassette's intrinsic capacity for coding information is ample, and the binding constraint is how often it fires, and not how many outcomes it can write. This is the quantitative reason the reconstructions in Figures S14 and S15 reach only a depth 9–10 and is why many clones are unresolved stars.

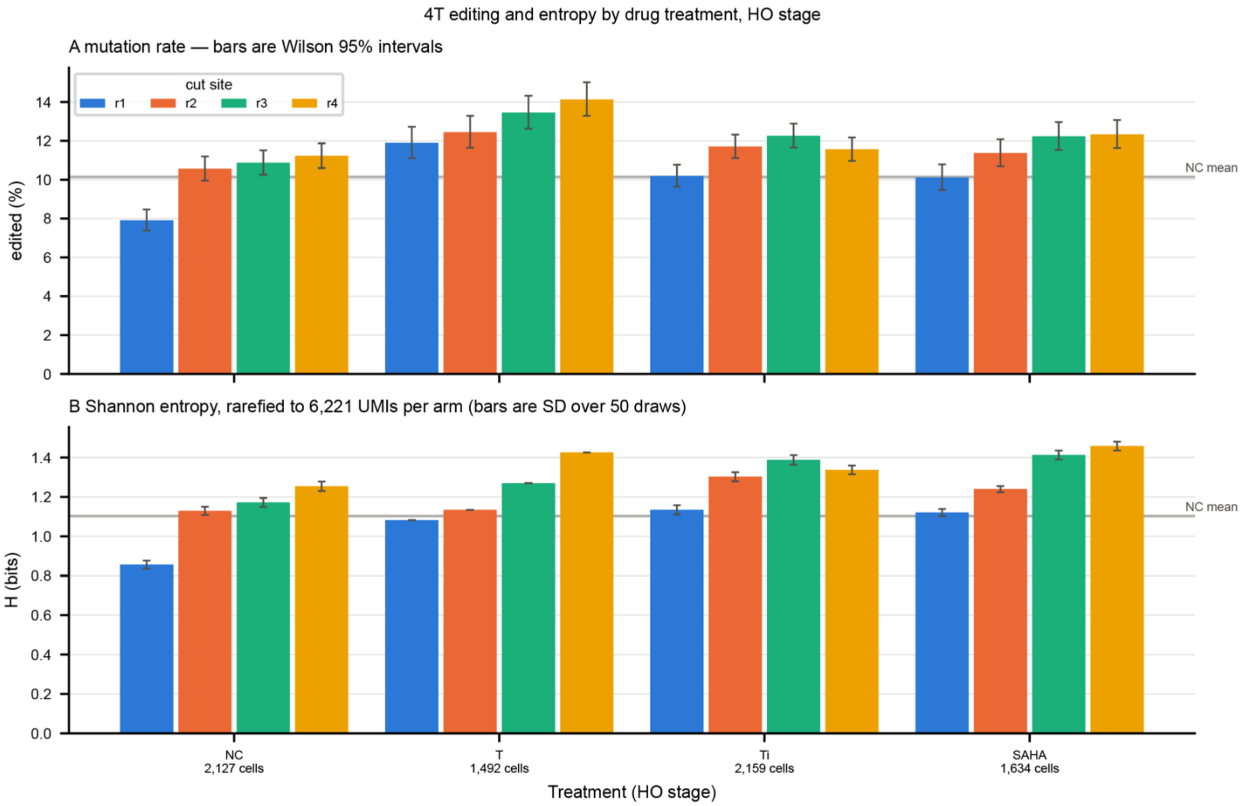


**Figure S13.** Cas12a editing and recorded allele diversity at the four 4 target cut sites by treatment arm at the mature organoid stage. microHOs were from the four arms compared in this study: NC: untreated control, T: TGFβ1 alone, Ti: TGFβ1 + SB431542, SAHA: TGFβ1 + SAHA. The data was pooled from four collection days (Day 20, 21, 26, 30) and four experimental batches,  with 492 to 2,159 cells and 6,221 to10,842 UMIs per arm. The four cut sites r1–r4 are color coded. (**A**) The percentage of UMIs carrying an indel at each cut site. An allele is called edited when Cassiopeia writes anything other than the original allele at that site, so a compound indel spanning one site's window still counts once. Error bars are Wilson 95% intervals on the binomial proportion (740 to 1,329 edited UMIs per arm × site). (**B**) Shannon entropy of the allele distribution at each cut site, in bits, rarefied to 6,221 UMIs per arm, error bars are the SD across draws. Even every treated arm showed a higher per-site edit rate than the untreated control, the cassette is delivered once before Day 0 and each site is cut once and destructively, so no treatment can create integrations or re-edit a site that has already fired. A higher edit rate under TGFβ1, with or without SAHA, is therefore consistent with either prolonged Cas12a activity in those arms or selective loss of cells whose sites were never cut; this design does not separate the two, and the quantity should not be read as a measure of how much lineage information the drug added.

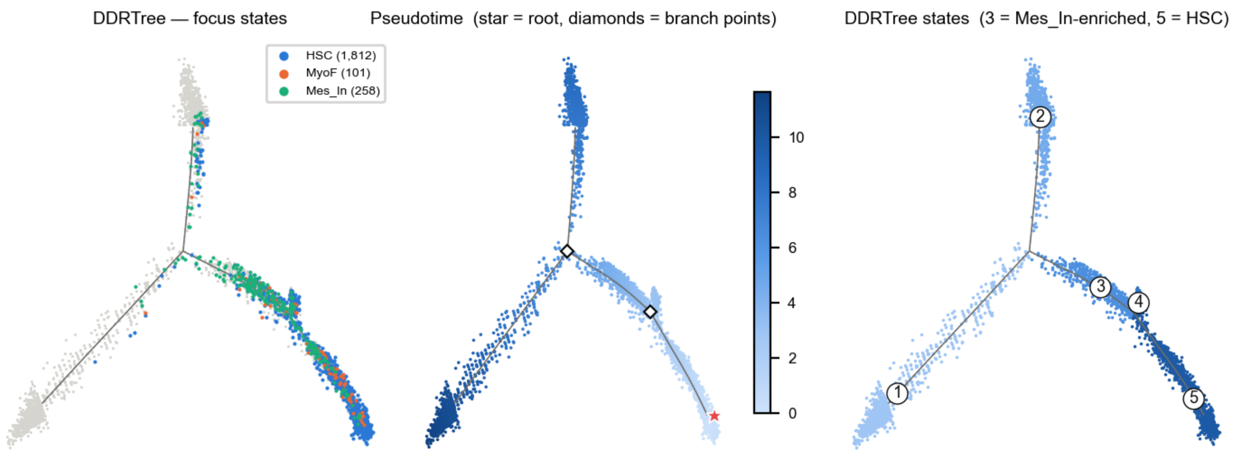


**Figure S14.** The DDRTrees drawn from the fitted geometry for 4,992 cells, 130 principal-graph points, and 129 edges. *Left:* The three focus cell states (HSC, MyoF and $Mes^{Inh}$) in color with every other cell shown in grey. *Middle:* Monocle 2 pseudotime graph is shown with the root cell (star) and two branch points (diamonds). The root cells was set to the state with the highest CytoTRACE2 potency, so the direction is fixed by an independent measurement. *Right:* The five DDRTree states with their numbers shown at the centroids. State 3 is the $Mes^{Inh}$-enriched branch and state 5 is the HSC branch, which has 76% HSCs in the tree. The grey skeleton shows the principal graph.
